# RIM-Binding Protein 2 organizes Ca^2+^ channel topography and regulates release probability and vesicle replenishment at a fast central synapse

**DOI:** 10.1101/2021.03.20.435607

**Authors:** Tanvi Butola, Theocharis Alvanos, Anika Hintze, Peter Koppensteiner, David Kleindienst, Ryuichi Shigemoto, Carolin Wichmann, Tobias Moser

## Abstract

RIM-Binding Protein 2 (RIM-BP2) is a multi-domain protein of the presynaptic active zone (AZ). By binding to Rab-interacting protein (RIM), bassoon and voltage-gated Ca²⁺ channels (Ca_V_), it is considered to be a central organizer of the topography of Ca_V_ and release sites of synaptic vesicles (SVs) at the AZ. Here, we investigated the role of RIM-BP2 at the endbulb of Held synapse of auditory nerve fibers with bushy cells of the cochlear nucleus, a fast relay of the auditory pathway with high release probability. Disruption of RIM-BP2 lowered release probability altering short-term plasticity and reduced evoked excitatory postsynaptic currents (EPSCs). Analysis of SV pool dynamics during high frequency train stimulation indicated a reduction of SVs with high release probability but an overall normal size of the readily releasable SV pool (RRP). The Ca^2+^-dependent fast component of SV replenishment after RRP depletion was slowed. Ultrastructural analysis by super-resolution light and electron microscopy revealed an impaired topography of presynaptic Ca_V_ and a reduction of docked and membrane-proximal SVs at the AZ. We conclude that RIM-BP2 organizes the topography of Ca_V_, and promotes SV tethering and docking. This way RIM-BP2 is critical for establishing a high initial release probability as required to reliably signal sound onset information that we found to be degraded in bushy cells of RIM-BP2-deficient mice *in vivo*.

**Significance Statement:** RIM-binding proteins (RIM-BPs) are key organizers of the active zone (AZ). Using a multidisciplinary approach to the calyceal endbulb of Held synapse that transmit auditory information at rates of up to hundreds of Hertz with sub-millisecond precision we demonstrate a requirement for RIM-BP2 for normal auditory signaling. Endbulb synapses lacking RIM-BP2 show a reduced release probability despite normal whole-terminal Ca^2+^ influx and abundance of the key priming protein Munc13-1, a reduced rate of SV replenishment, as well as an altered topography of Ca_V_2.1 Ca^2+^ channels, and fewer docked and membrane proximal synaptic vesicles. This hampers transmission of sound onset information likely affecting downstream neural computations such as of sound localization.

## Introduction

Active zones (AZs) are specialized regions at the presynaptic terminals where neurotransmitter release occurs. AZs employ a sophisticated machinery to enable ultrafast coupling of the incoming action potential to the release of transmitter via Ca^2+^-triggered SV fusion. Voltage-gated Ca^2+^ channels (Ca_V_) and SV release sites represent the core machinery, and their relative topography at the AZ co-determines the release probability (recent reviews in: Kaeser and Regehr, 2014; Schneggenburger and Rosenmund, 2015; Walter et al., 2018; Dittman and Ryan, 2019). The function and abundance of Ca_V_ (recent reviews in: Pangrsic et al., 2018; Dolphin and Lee, 2020) is positively regulated by auxiliary subunits and multi-domain proteins of the AZ such as RIM-BP, RIM, piccolo, bassoon, CAST and ELKS.

Several of these proteins promote the clustering of Ca^2+^ channels at the AZ and/or their interaction with the SV release sites (Gundelfinger and Fejtova, 2012; Südhof, 2012; Moser et al., 2019). RIM and RIM-BPs, in particular, have been considered as molecular linkers of Ca_V_ and vesicular release sites. Direct and indirect interactions of RIMs with Ca_V_ are well established (Kiyonaka et al., 2007; Gebhart et al., 2010; Kaeser et al., 2011; Picher et al., 2017). Proline-rich sequences of RIMs have been shown to bind the SH3-domains of RIM-BP that directly interacts with Ca_V_ (Hibino et al., 2002; Kaeser et al., 2011). Deletion of *Drosophila* RIM-BP disrupted Ca_V_ clustering at the AZs of larval neuromuscular junctions consequently impairing their functional coupling to SV release: altered short-term plasticity demonstrating a reduced release probability (Liu et al., 2011). Of the three mammalian RIM-BP isoforms, RIM-BP1 and 2 are neuron-specific, whereby RIM-BP2 seems to be the isoform that is most relevant for transmission at conventional synapses (Acuna et al., 2015; Grauel et al., 2016). Disruption of RIM-BP1 and 2 did not alter the Ca^2+^ current at the calyx of Held synapse (Acuna et al., 2015), while lack of RIM-BP2 (Krinner et al., 2017; Luo et al., 2017) and of RIM-BP1 (Luo et al., 2017) reduced the Ca_V_ number at ribbon-type AZs. An alteration of the Ca_V_ topography at the AZs has been reported based on super-resolution immunofluorescence microscopy (Grauel et al., 2016; Krinner et al., 2017).

A loosening of the otherwise tight spatial relationship of Ca_V_ and SV release sites was indicated by the reduced release probability and a greater sensitivity to the intracellular presence of the “slow” Ca^2+^ chelator EGTA in the absence of RIM-BP(1)/2 (Acuna et al., 2015; Grauel et al., 2016; Luo et al., 2017). A similar conclusion was reached in sensory hair cells where, unlike in CNS synapses, SV replenishment was impaired in the absence of RIM-BP2 (Krinner et al., 2017). However, in contrast to the dramatic impairment of synaptic transmission at RIM-BP deficient neuromuscular junctions in *Drosophila* (Liu et al., 2011), transmission was affected more subtly by RIM-BP deletion at mammalian synapses (Acuna et al., 2015; Grauel et al., 2016; Krinner et al., 2017; Luo et al., 2017). This might relate to partial compensation by other candidate linkers of Ca_V_ and SV release sites such as long RIM isoforms (Acuna *et al*, 2016). Recently, two alternative actions of RIM-BPs were indicated based on molecular perturbations studies in hippocampal neurons: i) binding to Ca_V_ enabling their tight coupling to SVs or ii) promoting SV priming by interaction with Munc13-1 (Brockmann et al., 2020). Here we aimed to further characterize the function of RIM-BP2 at the endbulb of Held synapse, the first central auditory synapse (von Gersdorff and Borst, 2002; Yu and Goodrich, 2014), that employs high vesicular release probability at its more than 100 AZs for reliable and temporally precise signal transmission from auditory nerve fibers (ANFs) to bushy cells (BCs) at frequencies of hundreds of Hertz (Trussell, 1999; Wang et al., 2011). We combined electrophysiological analysis with studies of the molecular composition and ultrastructure of the AZ in RIM-BP2-deficient endbulbs.

## Materials and Methods

### Animals

The constitutive RIM-BP2 knockout (RIM-BP2 KO) mice were described in (Grauel et al., 2016) and kindly provided to us by Drs. Christian Rosenmund, Katharina Grauel and Stephan Sigrist. All experiments were performed in compliance with the guidelines of the German animal welfare act and were approved by the board for animal welfare of the University Medical Center Göttingen and the animal welfare office of the state of Lower Saxony.

### Slice electrophysiology

Slice preparation: Acute parasagittal slices (150µm) from the anteroventral cochlear nucleus (aVCN) were obtained as described previously (Mendoza Schulz et al., 2014). Briefly, after sacrifice by decapitation, brains were dissected out and quickly immersed in ice-cold low Na^+^ and low Ca^2+^ cutting solution containing (in mM): 50 NaCl, 26 NaHCO_3_, 120 sucrose, 1.25 NaH_2_PO_4_.H_2_O, 2.5 KCl, 20 glucose, CaCl_2_, 6 MgCl_2_, 0.7 Na L-ascorbate, 2 Na pyruvate, 3 myo-inositol, 3 Na L-lactate with pH adjusted to 7.4 and osmolarity of around 310 mOsm/l. After removal of the meninges from the ventral face of the brainstem, the two hemispheres were separated by a midsagittal cut and the forebrain was removed at the pons-midbrain junction. The brain blocks containing brain stem and cerebellum were then glued (cyanoacrylate glue; Loctite 401, Henkel) to the stage of a VT 1200S vibratome (Leica microsystems, Wetzlar, Germany) such that the medial side was glued on, the ventral side was facing the blade and the lateral side was facing upwards, submerged in ice-cold cutting solution. For sectioning, the blade was positioned at the height of cerebellar flocculus and sections were cut at a blade feed rate of 0.02 mm/s with an amplitude of 1.50 mm. Slices were incubated for 30min in artificial cerebrospinal fluid (aCSF) maintained at 35°C, and then kept at room temperature (22-24°C) until recording. Composition of aCSF was identical to the cutting solution except (in mM): 125 NaCl, 13 glucose, 2 CaCl_2_ and 1 MgCl_2_. The pH of the solution was adjusted to 7.4 and osmolarity was around 310 mOsm/l. All solutions were continuously aerated with carbogen (95% O_2_, 5% CO_2_). For presynaptic endbulb recordings, coronal sections were made instead of parasagittal as described above. The only difference was that after the forebrain was removed at the pons-midbrain junction, the dissected brain was not cut midsagitally and the brain block containing the brain stem and the cerebellum was then glued to the vibratome stage with the caudal aspect facing upwards and the ventral side towards the blade.

Electrophysiology: Patch-clamp recordings were made from BCs of aVCN using EPC10 USB Patch clamp amplifier controlled by the Patchmaster software (HEKA Elektronik, Lambrecht/Pfalz, Germany). Sampling interval and filter settings were 25 µs and 7.3 kHz respectively. Cells were visualized by differential interference contrast (DIC) microscopy through a 40x water-immersion objective (NA 0.8; Zeiss, Oberkochen, Germany) using an Axioscope 2 FS plus microscope (Zeiss). All experiments were conducted at a temperature of 33-35°C, maintained by constant superfusion (flow rate 3-4 ml/min) of aCSF, heated by an inline solution heater (SH-27B with TC-324B controller; Warner Instruments, Hamden, CT, USA) and monitored by a thermistor placed between the inflow site and the slice, in the recording chamber.

Patch pipettes were pulled with P-87 micropipette puller (Sutter Instruments Co., Novato, CA, USA) from borosilicate glass capillaries with filament (GB150F, 0.86×1.50×80mm; Science Products, Hofheim, Germany). Open tip pipette resistance was 1.5-3 MΩ when filled with intracellular solution containing (in mM): 115 K-gluconate, 10 HEPES, 8 EGTA, 10 Na_2_Phosphocreatine, 4 ATP-Mg, 0.3 GTP-Na, 4.5 MgCl_2_, 10 NaCl and 1 *N*-(2,6-dimethylphenyl carbamoylmethyl) triethylammonium chloride (QX-314; Alomone Labs, Jerusalem, Israel) to block sodium channels, with a pH of 7.35 and an osmolarity of 300 mOsm/l. Additionally, 1 mM of fluorescent dye Alexa-488 (Invitrogen) was added to the recording pipette and cell structure was examined during experiments using a HXP 120 mercury lamp, with FITC filter (Semrock hardcoat). Cells were voltage-clamped at a holding potential of –70 mV, after correction for a liquid junction potential of 12 mV. Mean series resistance was around 5 MΩ and was compensated up to 70% with a 10 µs lag. Presynaptic ANF were minimally stimulated with a monopolar electrode in a patch pipette filled with aCSF, placed at a distance of at least three cell diameters from the cell being recorded. Stimulating currents of 10-20 µA were delivered through a stimulus isolator (A360 World Precision Instruments, Sarasota, FL, USA). For the main set of recordings, bath solution (aCSF) was supplemented with: 1 mM kynurenic acid sodium salt (abcam Biochemicals, Cambride, UK), a low-affinity AMPAR antagonist, to prevent receptor saturation/desensitization, 100µM Cyclothiazide to prevent AMPAR desensitization, 10 µM Bicuculline methchloride, a GABA_A_ receptor antagonist and 2 µM Strychnine hydrochloride, a glycine receptor antagonist.

For patch-clamp experiments at the endbulbs of Held patch-pipettes were coated with either dental wax or Sylgard to minimize fast capacitive transients and stray capacitance during voltage clamp experiments. Open tip pipette resistance was 4-5 MΩ with an intracellular solution containing (in mM): 130 Cs-methanesulfonate, 20 TEA-Cl, 10 HEPES, 0.5 EGTA, 5 Na_2_Phosphocreatine, 4 ATP-Mg, 0.2 GTP-Na, with pH adjusted to 7.3 with CsOH osmolarity of 320 mOsm/l. For anatomical confirmation 1 mM of fluorescent dye Alexa-488 (Invitrogen) was added to the recording pipette. The bath solution differed from the aCSF normally used (in mM): 85 NaCl, 25 glucose, 2 CaCl_2_ and 1 MgCl_2_. The pH of the solution was adjusted to 7.4 and osmolarity was around 310mOsm/l. Additionally, the bath solution was supplemented with 1 µM TTX, 1 mM 4-AP, and 40 mM TEA-Cl to suppress voltage-gated Na^+^ and K^+^ currents. Presynaptic terminals were voltage-clamped at a holding potential of –80 mV, a liquid junction potential of 3 mV was ignored. Series resistance was <30 MΩ and was compensated up to 50% with a 10 µs lag.

### Systems physiology: Extracellular recordings from single bushy cells

Extracellular recordings from single units of bushy cells (BCs) in the aVCN were performed as described before (Jing et al., 2013; Strenzke et al., 2016) on 9 to 10-week-old mice. After anesthetizing the mice with i.p. injection of urethane (1.32 mg/kg), xylazine (5 mg/kg) and buprenorphine (0.1 mg/kg), a tracheostomy was performed, their cartilaginous ear canals were removed and then they were positioned in a custom-designed head-holder and stereotactic system. After partial removal of the occipital bone and cerebellum to expose the surface of the cochlear nucleus, a glass microelectrode was advanced through the anterior portion of the aVCN to avoid the auditory nerve fibers and instead target the area with a higher fraction of spherical BCs. Acoustic stimulation was provided by an open field Avisoft ScanSpeak Ultrasonic Speaker (Avisoft Bioacoustics). “Putative” spherical BCs were identified and differentiated from other cell types in the cochlear nucleus by their characteristic ‘Primary-like’ peristimulus time histogram (PSTH) (Taberner and Liberman, 2005), irregular firing pattern demonstrated by a ≥ 0.5 coefficient of variation of inter-spike intervals of adapted responses, and a first spike latency of ≤ 5 ms. Bushy cells units were distinguished from auditory nerve fiber (ANFs, also having ‘Primary-like’ PSTH) based on their stereotactic position (<1.1 mm below the surface of the cochlear nucleus). Recordings were performed using TDT system III hardware and an ELC-03XS amplifier (NPI electronics).

### Immunohistochemistry and confocal imaging

Mice at postnatal day 20-24 were deeply and terminally anesthetized with Xylazin (5 mg/kg) and Ketamin (10 mg/ml) in 0.9% saline and then transcardially perfused with 2% freshly prepared ice-cold paraformaldehyde with pH adjusted to 7.4. The fixed brain was then removed and brainstem was dissected with a coronal cut few millimetres nasal to the junction between occipital cortex and cerebellum. The brain block was washed overnight in 30% sucrose solution in PBS. For sectioning, the brain block was embedded in Tissue Tek Cryomatrix (Thermo Fisher Scientific, Waltham, MA, USA) and then fixed on the stage of the cryostat (Figocut E cryotome, Reichert-Jung, Depew, NY, USA) such that the caudal aspect was facing upwards and the dorsal side was towards the blade. Advancing from caudal to nasal, 30 µm coronal sections were cut (chamber temperature: −20°C, object temperature: −22°C) and discarded until the appearance of the 7^th^ cranial nerve. Subsequent sections containing aVCN were collected onto electrostatically charged microscope slides (SuperFrost Plus, ThermoFisher Scientific, MA, USA). For parallel processing, one slice of each genotype was collected per slide.

Thereafter, the slices were washed for 10 min in PBS and incubated in Goat Serum Dilution Buffer (GSDB; 16% normal goat serum, 450 mM NaCl, 0.3% Triton X-100, 20 mM phosphate buffer, pH 7.4) for 1 h, followed by incubation in primary antibodies diluted in GSDB, for 3 h, in a wet chamber at room temperature. After washing 2×10 min with wash buffer (450 mM NaCl, 0.3% Triton X-100, 20 mM phosphate buffer) and 2×10 min with PBS, the slices were incubated with secondary antibodies diluted in GSDB, for 1 h, in a light-protected wet chamber at room temperature. The slices were then washed 2×10 min with wash buffer, 2×10 min with PBS and 1×10 min in 5 mM phosphate buffer, and finally mounted with a drop of fluorescence mounting medium based on Mowiol 4-88 (Carl Roth, Karlsruhe, Germany) and covered with a thin glass coverslip. The above described perfusion fixation method was used to stain for RIM-BP2 (Figure 4 A, B). The remaining immunofluorescence experiments (Munc13-1, RIM2, CAST, bassoon) were done on samples taken from RIM-BP2 KO mice aged between p15 – p23 and WT littermates. After obtaining coronal sections from the cryostat the slices were maintained frozen at −20°C until immersion fixation or fixed directly on ice in a solution of PBS containing 3% w/v heat depolymerized PFA (70°C) for 3 minutes. This alternative method of fixation showed robust labelling with less background than perfusion fixed samples labelled against the same markers. For the comparison of Ca_V_2.1 immunofluorescence levels in confocal microscopy and the STED analysis of Ca_V_2.1 and bassoon clusters, we used very brief fixation with 3% PFA for less than one minute (30-40 seconds) as Ca_V_2.1 labelling was impeded by stronger fixation. Thereafter, the blocking and immunolabelling protocols were followed exactly as described above for the perfusion fixed samples.

**Figure 4.**
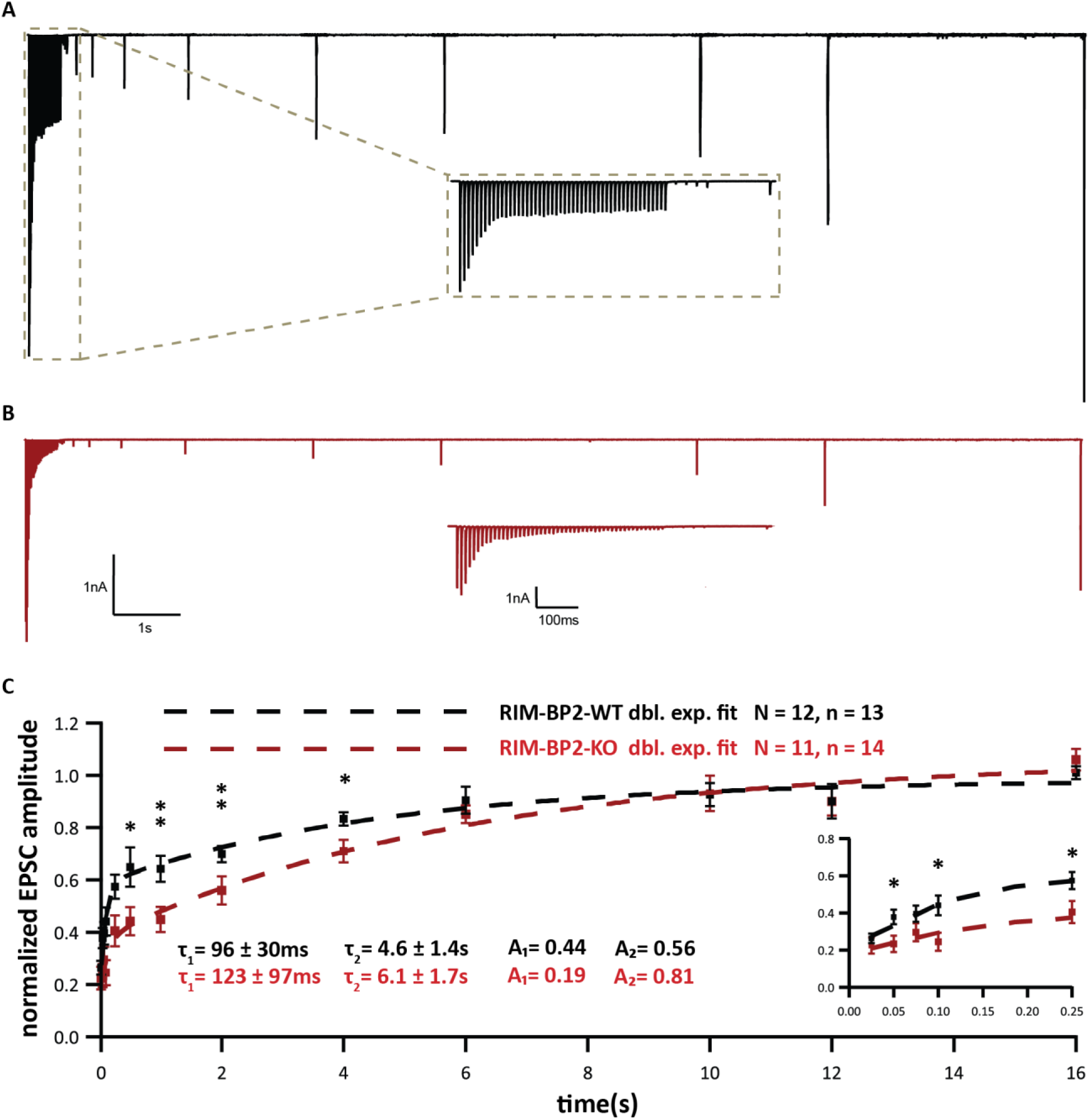
Slowed RRP recovery in RIM-BP2 deficient endbulb synapses. (**A-B**) Representative traces of RIM-BP2 WT (**A**) and RIM-BP2 KO (**B**), illustrate the recovery experiment. After a 100 Hz conditioning train of 50 stimuli, single test pulses were delivered at time intervals of (in ms) 25, 50, 75, 100, 250, 500 (further intervals in s), 1, 2, 4, 6, 10, 12 and 16. To assess recovery, the EPSC amplitude in response to the test pulse is normalised to the first EPSC amplitude of the conditioning train. Insets (**A, B**) show the time course of recovery during the first 5 test stimuli in sub-second detail. (**C**) Recovery is plotted as mean ± SEM EPSC amplitude in response to test pulses normalized to the first EPSC amplitude of the conditioning train. The double exponential fits are represented by the dashed lines for RIM-BP2 WT (black) and RIM-BP2-KO (red). The time constants (τ) and fractional contributions (A) of fast (τ_1_, A_1_) and slow (τ_2_, A_2_) recovery components are provided on the graph. Inset shows the first five responses in detail. Normality was tested with Jarque-Bera test. Statistical significance between groups was tested with Mann-Whitney U-test. \*\**p-value* < 0.01, \**p-value* < 0.05. *N*: number of animals, *n*: number of BCs

Primary antibodies used were: rabbit anti-RIM-BP2 (1:200), guinea pig anti-VGLUT1 (1:500), rabbit anti-VGLUT1 (1:1000), mouse anti-gephyrin (1:500), mouse anti-Sap7f407 to bassoon (1:500; Abcam, Cambridge, UK), guinea pig anti-bassoon (1:500), rabbit anti-Munc13-1 (1:200), rabbit anti-RIM1 (1:200), rabbit anti-RIM2 (1:200), rabbit anti-CAST (1:200), rabbit anti-P/Q Ca^2+^ channel (1:500), chicken anti homer1 (1:200). Unless stated otherwise, primary antibodies were purchased from Synaptic Systems, Göttingen, Germany. Secondary antibodies used were: AlexaFluor488-, AlexaFluor568- and AlexaFluor647-labeled antibodies (1:200, Invitrogen), goat anti-guinea pig STAR580 and goat anti-rabbit STAR 635p (1:200, Abberior GmbH, Goettingen, Germany).

Confocal images were acquired using a laser-scanning confocal microscope (Leica TCS SP5; Leica Microsystems) equipped with 488 nm (Ar) and 561/633 nm (He-Ne) lasers and 63x/1.4 NA oil-immersion objective. STED and confocal images (Ca_V_2.1 and bassoon analyses) were acquired using a 2-color STED microscope (Abberior Instruments, Göttingen, Germany) equipped with 561 and 640 nm excitation lasers, a 775 nm laser for STED (1.2W) and a 100x oil immersion objective (1.4NA, Olympus). Confocal z-stacks were processed with Imaris (Bit-plane, Zurich, Switzerland) for spot detection, co-localization analysis and fluorescence intensity calculation using custom Matlab scripts. STED images of Ca_V_2.1 and bassoon were analyzed using Igor Pro7 (Wavemetrics, Lake Oswego, OR, USA). Samples of both genotypes: RIM-BP2 WT and RIM-BP2 KO were processed and imaged in parallel, using same laser power, gain and microscope settings.

### SDS – Freeze-fracture Replica Immuno-Labelling (SDS-FRIL)

Mice at postnatal day 20-24 were deeply and terminally anesthetized with Xylazin (5 mg/kg) and Ketamin (10 mg/ml) in 0.9% saline, and perfused transcardially with ice-cold PBS followed by perfusion with freshly-prepared 2% PFA with 15% saturated picric acid solution in 0.1 M phosphate buffer (PB) with pH adjusted to 7.3. The fixed brain was then removed and brainstem was dissected with a coronal cut few millimetres nasal to the junction between occipital cortex and cerebellum. The brain block was washed over night in 30% sucrose solution in PBS. Coronal slices (130 µm thick) were cut from the fixed brain block fusing a vibratome microslicer (Linear-Pro7, Dosaka) in ice-cold PBS. The rostral anterior ventral cochlear nuclei were trimmed by hand from the slices. The trimmed sections were then immersed in graded glycerol concentrations of 10%-20% at room temperature for 20 min each, followed by 30% at 4°C overnight. The trimmed sections were sandwiched between two metal carriers and then rapidly frozen by a high-pressure freezing machine (HPM010, BAL-TEC, Balzers). Using a freeze etching device (BAF060, BAL-TEC), frozen samples were then fractured into two parts at −115°C, and the fractured faces were replicated by sequential deposition of carbon (thickness: 5 nm from 90° angle), platinum (thickness: 2 nm from 60° angle, unidirectional), and carbon again (thickness: 20 nm from 90° angle). After thawing, the tissue debris attached to the replica was digested with gentle shaking at 80°C for 18 h, in a solution containing 2.5 % SDS, 20% sucrose, and 15 mM of Tris-HCl with pH set to 8.3. The replicas were washed 3 x 10 min in wash buffer (0.1% Tween-20, 0.05% BSA, 0.05% NaN_3_ in TBS, pH 7.4), and then the non-specific binding sites were blocked with 5% BSA in wash buffer for 1 h at 4°C.

For multiple immunolabelling against Ca_V_2.1 and AZ proteins, replicas were first incubated with guinea pig anti-Ca_V_2.1 antibody (8 µg/ml in 1% BSA; Frontier Institute AB_2571851) at 15°C for 3 days, then with anti-guinea pig secondary antibodies conjugated with 10 nm gold particles (1:30 diluted in 5% BSA; British Biocell International) at 15°C overnight, followed by incubation with a cocktail of rabbit anti-AZ proteins antibodies (anti-ELKS at 2 µg/ml in 1% BSA; gift from Prof. Ohtsuka raised against rat ELKS aa117-142, anti-Neurexin at 4 µg/ml in 1% BSA; gift from Prof. Watanabe raised against aa1499-1507 and anti-RIM at 4 µg/ml in 1% BSA; Synaptic Systems 140203) at 15°C overnight, and finally with anti-rabbit secondary antibodies conjugated with 5 nm gold particles (1:30 diluted in 5% BSA; British Biocell International) at 15°C overnight. After immunolabelling, replicas were rinsed three times with 0.05% BSA in TBS, washed with TBS and distilled water, and mounted on formvar-coated copper grids.

The labelled replicas were imaged using a Tecnai-12 transmission electron microscope (FEI; AV 120 kV). To obtain a planar view for quantitative measurement of immuno-gold particle number and densities, profile of synaptic structures (AZ and PSD) were tilted in the electron beam. IMP clusters representing PSDs were manually demarcated and the area was measured using ImageJ software (Rubio et al., 2017). Active zone areas were marked by hand with the experimenter being blinded to the identity of the two genotypes. Quantitative analysis of immune-gold particles was done using an in-house software tool – Gold Particle Detection and Quantification (Luján et al., 2018). To define clusters of gold particles the threshold for the distance between particles belonging to the same cluster was calculated as µ + 2σ, where µ and σ are the mean and standard deviation obtained from a Gaussian fit to the distribution of nearest neighbour distances (NND) between particles. µ + 2σ was 43.02 nm and 43.92 nm for Ca_V_2.1 gold particles in RIM-BP2 WT and KO respectively. We set the threshold at 40 nm to match the value used in a previous analysis (Miki et al., 2017). The distances between particles were measured from their centers of mass, and the minimum number of particles required to form a cluster was set to three. We additionally compared our ‘real’ distribution of NNDs between gold particles and their clustering to 500 random and fitted simulations (keeping the NNDs between the simulated particles similar to the ones between the ‘real’ gold particles) generated by Monte-Carlo simulations (as described in (Luján et al., 2018; Kleindienst et al., 2020)) to confirm that the clusters visualized through our analysis are not generated by chance.

### High-pressure freezing, Freeze substitution and Electron tomography

Parasagittal slices from cochlear nuclei were obtained as described for slice electrophysiology. Slices containing the cochlear nucleus were trimmed and mounted onto type A specimen carriers (Leica Microsystems, Wetzlar, Germany) filled with cutting solution. The flat side of the type B carriers (Leica Microsystems, Wetzlar, Germany) was dipped in 1-hexadecene (Sigma-Aldrich, Wetzlar, Germany) and placed onto the type A carriers. Samples were frozen immediately using a HPM100 (Leica Microsystems, Wetzlar, Germany) and transferred into liquid nitrogen. Freeze-substitution was performed in an EM AFS2 (Leica Microsystems, Wetzlar, Germany) according to Wong *et al*. (Wong et al., 2014). The slices were incubated in 0.1% (w/v) tannic acid in acetone at −90°C for 4 days and afterward washed three times for 1 h each in acetone at −90°C. 2% (w/v) osmium tetroxide in acetone was applied and incubated for 40.4 h. During that time the temperature was raised slowly to 4°C (10°C/h). At 4°C, osmium tetroxide was removed, and the samples were washed with acetone three times and brought to room temperature. Slices were infiltrated in epoxy resin (Agar-100 kit, Plano, Germany; epoxy/acetone 1:1 3-6 h; 100% epoxy overnight). Finally, samples were further incubated in fresh 100% epoxy and placed in embedding molds.

After polymerization for 48 h at 70°C, excess resin was removed with a fine file (DiAtome, Switzerland) and the block was trimmed to a pyramid using a razor blade. To check the region and the structural preservation, 65 nm ultrathin sections were cut with a diamond knife (DiAtome, Switzerland) using an EM UC7 (Leica Microsystems, Wetzlar, Germany) ultramicrotome. Sections were collected on formvar-coated copper slot grids (Athene, Plano, Wetzlar, Germany, for ultrathin sections). For electron tomography, 250 nm semi-thin sections were obtained and collected on mesh grids (100 mesh; Athene, Plano, Wetzlar, Germany, for semi-thin sections). Post-staining was performed with Uranyless (EMS, Hatfield, PA) for 20 min.

The region and quality of the tissue was checked at 80 kV using a JEM1011 transmission electron microscope (JEOL, Freising, Germany) equipped with a GatanOrius 1200A camera (Gatan, Munich, Germany). Electron tomography was performed as described previously (Wong et al., 2014). 10 nm gold beads (British Bio Cell/Plano, Germany) were applied to both sides of the stained grids. Big synaptic terminals on bushy cells were identified and tilt series from endbulb AZs were acquired at 200 kV using a JEM2100 transmission electron microscope (JEOL, Freising, Germany) mostly from −60° to +60° with a 1° increment at 15,000× using the Serial-EM software package with an image pixel size of 0.95 nm (Mastronarde, 2005). Tomograms were generated using the IMOD package etomo (Kremer et al., 1996).

Only asymmetric synapses with clearly identifiable PSDs were analyzed. However, in high-pressure frozen samples, PSDs appear less electron-dense compared to chemical fixed synapses. Only AZs that showed a PSD and a clear synaptic cleft, originating from large presynaptic terminals were analyzed to exclude inhibitory synapses.

Tomograms were segmented semi-automatically using 3dmod (Kremer et al., 1996). The AZ membrane was manually segmented every 15 virtual sections for five consecutive virtual sections and then interpolated across the Z-stack, following the extent of the PSD and the parallel synaptic cleft. Moreover, virtual sections were corrected manually after interpolation. The total surface area of this object was then divided by two to calculate the AZ area.

Synaptic vesicles (SVs) were reconstructed at their maximum projection and the sphere size was adjusted for each vesicle. The smallest distances from the outer leaflet of the SV membrane to the inner leaflet of the AZ membrane were measured and SVs in contact with the AZ membrane were defined as morphological docked SVs (0-2 nm distance). Moreover, all vesicles within 200 nm of the AZ were quantified and categorized in 20 nm bins. The radii of the SVs were determined with the program “imodinfo” of the IMOD software package and the diameters were calculated. For quantification of lateral distances of docked SVs, models of tomogram top-views with docked SVs were imported in ImageJ and the center of the captured AZ area was defined by setting two diagonal lines from respective edges of the AZ. The crossing point was defined as the center and the distances were measured from the outer membrane of the SV towards the center point.

### Data analysis

Electrophysiology data were analyzed using Igor Pro (Wavemetrics, Lake Oswego, OR, USA), Mini Analysis (Synaptosoft Inc., Fort Lee, NJ, USA) and GraphPad Prism software (La Jolla, CA, USA). Synaptic delay was calculated as the time between the start of stimulus (voltage output of the amplifier as dictated by the experiment protocol) and the time when the respective EPSC response reached 10% of its peak amplitude.

Confocal images were analyzed using ImageJ software, Imaris (Bitplane AG, Zurich, Switzerland) and Matlab (Mathworks). STED images of Ca_V_2.1 channels were analysed using custom Igor pro scripts. Single plane slices of the top (coverslip proximal) and/or bottom (distal from coverslip) membranes of BCs were imaged to capture the Ca_V_2.1 spots on a flat surface top view and enable 2D Gaussian fitting. The Ca_V_2.1 spots that were simultaneously closely juxtaposed to puncta of both homer1 and bassoon were fitted with Igor Pro’s 2D Gaussian function. Figures were assembled for display using Adobe Illustrator (Adobe Systems, Munich, Germany). Unless reported otherwise, statistical significance between groups was determined by either unpaired Student’s t-test (in case of normally distributed data with comparable variances between the groups) or Wilcoxon rank sum test (when data distribution did not satisfy the criteria). Normality of distribution was tested with Jarque-Bera test and variances were compared with F-test. Data were presented as mean ± S.E.M. when compared using Student’s t-test. In case of Wilcoxon’s rank sum test, data were presented as box and whisker plots showing grand median (of the means of all recordings), lower/upper quartiles, 10-90^th^ percentiles). *, **, ***, **** indicate p < 0.05, 0.01, 0.001 and 0.0001 respectively.

## Results

### Deletion of RIM-BP2 impairs synchronous transmitter release at the endbulb of Held synapse

To determine the functional role of RIM-BP2, we studied synaptic transmission at the endbulb of Held synapse in acute parasagittal slices of the brainstem of constitutive RIM-BP2 knockout mice (RIM-BP2 KO, (Grauel et al., 2016)) recording spontaneous (no stimulation, no TTX applied) and evoked excitatory postsynaptic currents (sEPSCs and eEPSCs, respectively) from bushy cells (BCs) of the anteroventral cochlear nucleus (aVCN, Fig. 1A) at postnatal days 15-21. eEPSCs were elicited by minimal electrical stimulation of the presynaptic ANF by a monopolar electrode placed in the proximity (∼3 cell diameters away) of the recorded BC, whereby each stimulus is aimed to elicit one action potential in one endbulb (Yang and Xu-Friedman, 2008).

**Figure 1.**
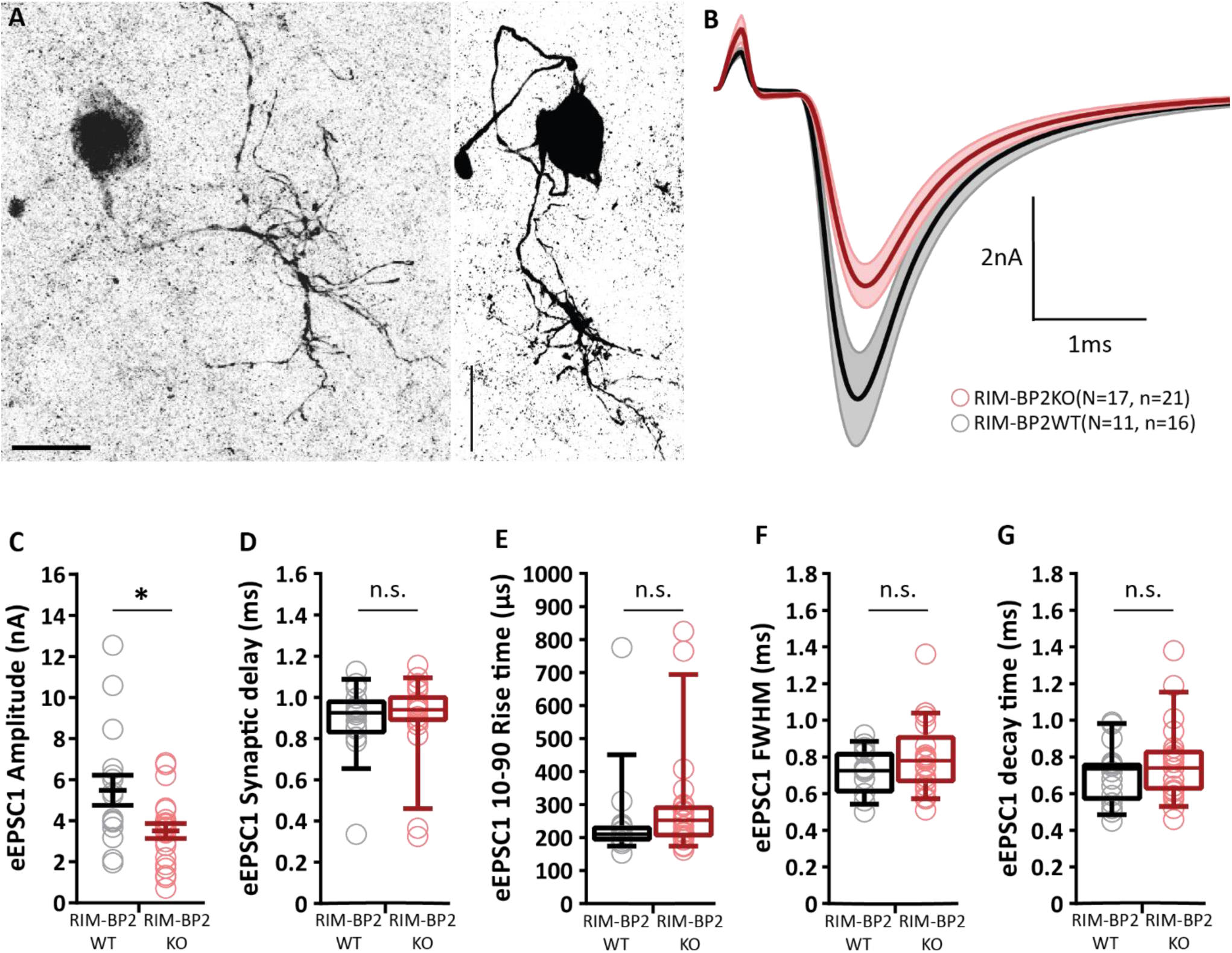
Reduced amplitude of evoked EPSCs in RIM-BP2 deficient endbulb synapses. (**A**) BCs (left and right panels showing dye-filled BCs) were distinguished from stellate cells (another major cell type in the aVCN) by the faster kinetics of their postsynaptic currents (Isaacson and Walmsley, 1995) and their characteristic short-term plasticity (Chanda and Xu-Friedman, 2010). In addition to such functional identification, each recorded cell was filled with fluorescent dye Alexa 488 via the patch pipette for morphological distinction. BCs are spherical in appearance with one primary dendrite terminating in a dense bush-like dendritic tree (Wu and Oertel, 1984), distinct from stellate cells, which are asymmetrical in shape and have multiple dendrites branching off in various directions giving them a star-like appearance. Scale bars: 20 µm. (**B**) Average traces (± SEM) of eEPSCs in RIM-BP2 KO (red) and WT (black) endbulbs. (**C**) Reduced eEPSC amplitude in RIM-BP2 KO compared to WT. (**D-G)** No significant differences in eEPSC kinetics: (**D**) synaptic delay, (**E**) 10-90% rise time, (**F**) full width at half-maximum and (**G**) decay time. Data points represent the mean estimate of each BC included in the analysis. Normality was tested with the Jarque-Bera test. Normally distributed data (**C**) are shown as grand mean (of the means of all BCs) ± SEM. Non-normally distributed data (**D, E, F, G**) are presented as box and whisker plots (grand median of all BC means, lower/upper quartiles, 10-90^th^ percentiles); n.s. *p-value* ≥ 0.05, Mann-Whitney U-test. *N*: number of animals, *n*: number of BCs.

Experiments of Figure 1 were performed in the presence of 1 mM kynurenic acid and 100 µM Cyclothiazide (CTZ) to avoid saturation and desensitization of AMPA receptors (Chanda and Xu-Friedman, 2010), respectively. The concentration of CTZ used here, was adopted from previous reports at the calyx of Held (Sakaba and Neher, 2001; Thanawala and Regehr, 2013). The former study (Sakaba & Neher, 2001) examined the NMDA current in the presence of 100 µM CTZ and found no evidence for a presynaptic CTZ effect at the calyx of Held. The eEPSC amplitude was reduced in RIM-BP2 KO BCs as compared to littermate wildtype (WT, Fig. 1B-C, 3.50 ± 0.37 nA (KO) vs. 5.48 ± 0.73 nA (WT), *p* = 0.022, Mann-Whitney U-test). There was a tendency towards slower eEPSC kinetics, but this did not reach significance for either synaptic delay (Fig. 1D) or rise and decay time (Fig. 1E-G), (p-value ≥ 0.05, Mann-Whitney U-test). EPSC recordings in the absence of kynurenic acid and CTZ (Fig. 1-1) confirmed the eEPSC reduction, eliminating the possibility of CTZ obscuring the changes in AMPA receptor composition or inducing membrane potential changes. The eEPSC kinetics was significantly slower for RIM-BP2 KO BCs under these conditions (Fig. 1-1).

**Figure 1-1.**
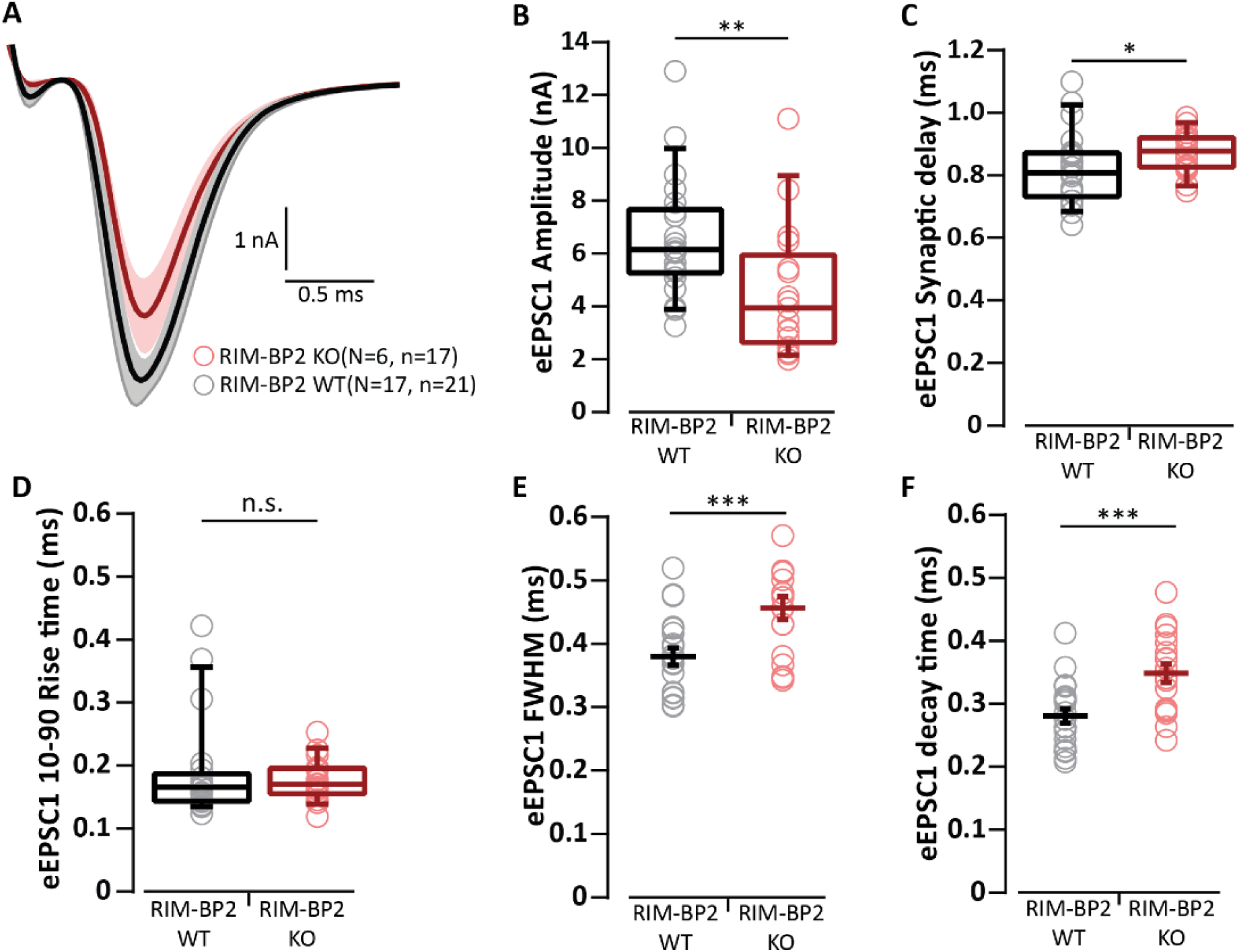
Evoked EPSC recordings in the absence of kynurenic acid and CTZ. (**A**) Average traces (± SEM) of eEPSCs in RIM-BP2-KO (red) endbulbs and WT (black) in the absence of kynurenic acid and CTZ. (**B**) Reduced eEPSC amplitude in RIM-BP2 KO compared to in WT. (**C-F**) eEPSC kinetics: (**C**) a larger synaptic delay, (**D**) unchanged 10-90% rise time, (**E**) wider full-width at half-maximum, and (**F**) slower decay time. Data points represent the mean estimate of each BC included in the analysis. Normality was tested with the Jarque-Bera test. Normally distributed data (**E, F**) are shown as grand mean (of the means of all BCs) ± SEM, and significance was tested with Student’s t-test. Non-normally distributed data (**B, C, D**) are presented as box and whisker plots (grand median of all BC means, lower/upper quartiles, 10-90^th^ percentiles), and significance was tested with Mann-Whitney U-test; n.s. *p-value* ≥ 0.05, * *p-value < 0.05, **p-value <0.01, *** p-value <0.001. N*: number of animals, *n*: number of BCs.

In order to test for potential changes in the quantal release properties we recorded spontaneous EPSCs (sEPSCs) from BCs (Fig. 1-2). We did not observe differences in the sEPSC amplitude (Fig. 1-2A-C), kinetics (Fig. 1-2D-F) and frequency (Fig. 1-2G; p-value ≥0.05 for all 3 quantities), which was also the case when recording in the absence of kynurenic acid and CTZ (Table 1). Unaltered sEPSCs suggest that the properties of single SV release and of postsynaptic glutamate response are intact in RIM-BP2 deficient endbulb synapses at BCs.

**Figure 1-2.**
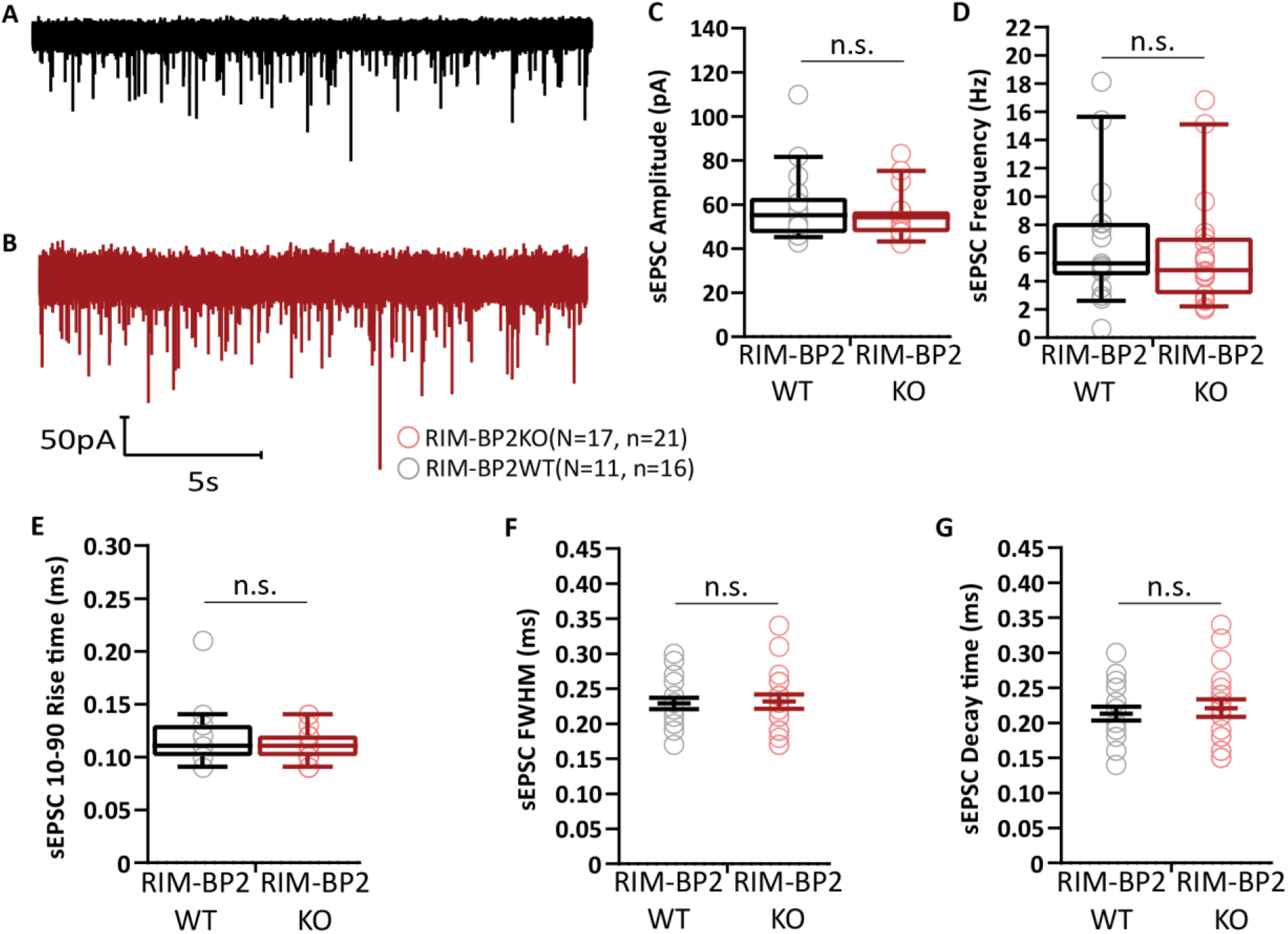
Unaltered amplitude, kinetics and frequency of spontaneous EPSCs in RIM-BP2 deficient endbulb synapses. (**A, B**) Representative sEPSCs at WT (**A**, black) and RIM-BP2 KO (**B**, red) endbulbs of held: continuous traces are shown. No significant differences in (**C**) amplitude, (**D**) frequency, (**E**) 10-90% rise time, (**F**) full-width at half-maxima and (**G**) decay time in RIM-BP2 deficient endbulbs. sEPSC were recorded while intracellularly blocking BC action potential generation with QX-314. Each data point represents the mean estimate of each BC included in the analysis. Normality was tested with the Jarque-Bera test. Normally distributed data are depicted as mean (grand average of all BC means) ± SEM (**F**, **G**; n.s. *p-value* ≥ 0.05, unpaired Student’s t-test). Non-normally distributed data are presented as box and whisker plots (grand median of all BC means, lower/upper quartiles, 10-90^th^ percentiles) (**C, D, E**); n.s. *p-value* ≥ 0.05, Mann-Whitney U-test *N*: number of animals, *n*: number of BCs.

**Table 1.** Unaltered sEPSC at the RIM-BP2 deficient endbulb of Held in the absence of kynurenic acid and CTZ. Data presented as mean (grand average of the means of all BCs) ± S.E.M. Statistical significance between groups was determined by either unpaired Student’s t-test (in case of normally distributed data with comparable variances between the groups) or Mann-Whitney U-test (when data distribution did not satisfy the criteria). Normality of distribution was tested with Jarque-Bera test and variances were compared with F-test. WT N = 4; n = 10, RIM-BP2 KO N = 7; n = 19 (N, number of animals; n, number of BCs).

| Parameter | WT | RIM-BP2 KO | p-value |
| --- | --- | --- | --- |
| Amplitude (pA) | 130.54 ± 12.06 | 112.48 ± 4.03 | 0.22 |
| 10-90% Rise time (ms) | 0.09 ± 0.003 | 0.09 ± 0.002 | 0.85 |
| FWHM (ms) | 0.18 ± 0.008 | 0.19 ± 0.004 | 0.46 |
| Decay time (ms) | 0.16 ± 0.010 | 0.17 ± 0.005 | 0.36 |
| Frequency (Hz) | 7.65 ± 1.28 | 8.01 ± 1.09 | 0.83 |

Quantal size being unaltered, the reduced eEPSC alteration could result from either: (i) an impaired stimulus-secretion coupling due to reduced Ca^2+^ influx and/or altered topography of Ca^2+^ channels (Krinner et al., 2017) and fusion competent SVs (Acuna et al., 2015; Grauel et al., 2016; Luo et al., 2017), or (ii) due to an impaired SV priming that has recently been shown to involve RIM-BP2 interaction with Munc13-1 (Brockmann 2020). We first checked for changes in presynaptic Ca^2+^ influx in the absence of RIM-BP2 using ruptured-patch recordings from the endbulb of Held (Lin et al., 2011) in mice after the hearing onset (postnatal days 13-16, Fig. 2). We did not observe significant changes in the peak Ca^2+^ current amplitude or peak Ca^2+^ current density in KO endbulbs of Held (Fig. 2).

**Figure 2.**
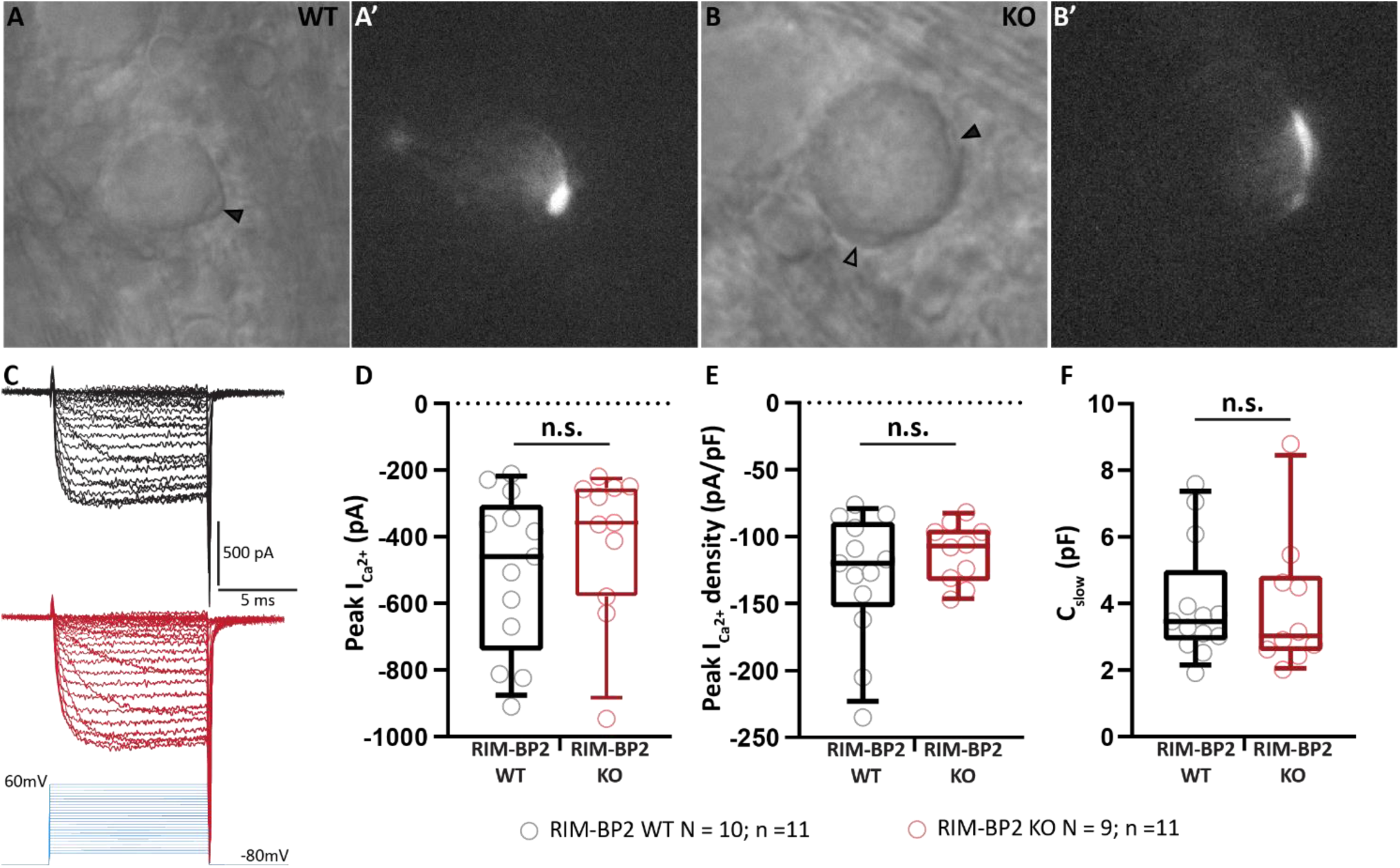
Ca^2+^ influx is unaltered in RIM-BP2 deficient endbulbs. (**A, B**) Bright-field images of the WT (**A**) and RIM-BP2 KO (**B**) bushy cells under DIC. The recorded endbulb is demarcated by a solid black arrow head (**A, B**), while another endbulb onto to the same bushy cell is shown by an open black arrowhead (**B**). Each recorded endbulb was filled with Alexa 488 to confirm that only the presynaptic terminal was accessed, and not the postsynaptic cell. Fluorescent endbulbs are shown in **A’, B’**. (**C**) Representative traces of presynaptic Ca^2+^ currents in WT (black) and RIM-BP2 KO (red) in response to stepped increase in holding potential (stimulus protocol in blue). (**D-F)** No significant difference was observed between the two genotypes in the peak Ca^2+^ current (**D**), peak Ca^2+^ density (**E**; peak current/C_slow_), and the size of the endbulb (**F**, estimated by C_slow_). Each data point represents the mean estimate of each endbulb included in the analysis. Normality was tested with the Shapiro-Wilk test. Non-normally distributed data are presented as box and whisker plots (grand median of all BC means, lower/upper quartiles, 10-90^th^ percentiles); n.s. *p-value* ≥ 0.05, Mann-Whitney U-test *N*: number of animals, *n*: number of endbulbs of Held.

Next, we investigated whether eEPSC alteration is caused by a reduction in release probability or altered vesicle pool dynamics. We used high frequency stimulation to assess short-term plasticity, release probability as well as the size and dynamics of the RRP during *quasi* physiological regimes of synaptic transmission. Fifty consecutive stimuli were delivered at 100, 200, and 333 Hz (Fig. 3: in the presence of 1 mM kynurenic acid and 100 µM CTZ; Fig. 3-1 in the absence of both drugs). Different from the prominent short-term depression typical for the endbulb of Held (Oleskevich and Walmsley, 2002; Yang and Xu-Friedman, 2008), RIM-BP2 deficient endbulb synapses showed an initial facilitation followed by slower depression (Fig. 3, Table 2, Fig. 3-1). Facilitation was evident from the paired pulse ratio (PPR) > 1 in RIM-BP2 deficient endbulb synapses for all inter-stimulus intervals tested, while PPR was consistently ∼ 0.8 in WT (Table 2). The extent of depression assessed as EPSC_30-50_/EPSC_max_ (Fig. 3, Table 2, Fig. 3-1) tended to be less in RIM-BP2 deficient endbulb synapses without reaching statistical significance. Release probability (P_r_) as well as the size and dynamics of the RRP were estimated by applying two variants of the cumulative analysis to the EPSC trains (Fig. 3): Schneggenburger-Meyer-Neher (SMN) method (Schneggenburger et al., 1999) and Elmqvist & Quastel (EQ) method (Elmqvist and Quastel, 1965); for review see: (Neher, 2015). Both methods revealed a significant reduction of P_r_ in RIM-BP2 KO synapses while RRP size and the replenishment rate were not significantly altered (Table 2). In an attempt to address the question whether the reduced release probability reflects a smaller complement of high release probability SVs (“tightly docked” or “super-primed” SVs (Neher and Brose, 2018)) we followed a previously described analysis that found a reduction of SV super-primed SVs upon genetic deletion of all rab3 isoforms (Schlüter et al., 2006). We subtracted the quantal content of each RIM-BP2 KO response from the respective RIM-BP2 WT response during train stimulation (Fig. 3M-O). This analysis revealed the strongest difference between the two genotypes occurs at the beginning of the train, where the RIM-BP2 KO endbulbs release 30-40 SVs less than the RIM-BP2 WT, which is consistent with a reduction of high release probability (“super-primed”) SVs in the absence of RIM-BP2. The difference vanishes already after the first two eEPSCs, when the subtraction curves for 200 and 333 Hz actually cross the zero line, indicating that release from KO-synapses is actually slightly larger than that from WT. Possible reasons for this small difference include i) protracted release of low release probability SVs that could result from impaired Ca_V_-release site coupling and ii) residual desensitization that is more prominent for RIM-BP2 WT synapses with larger eEPSCs (despite 100 µM cyclothiazide and 1 mM kynurenic acid). During the steady state response to the train stimulation, the WT synapses tended to release more SVs than the KO (average difference: 6 SVs for 100 Hz, 3 SVs for 200Hz, no difference for 333 Hz).

**Figure 3-1:**
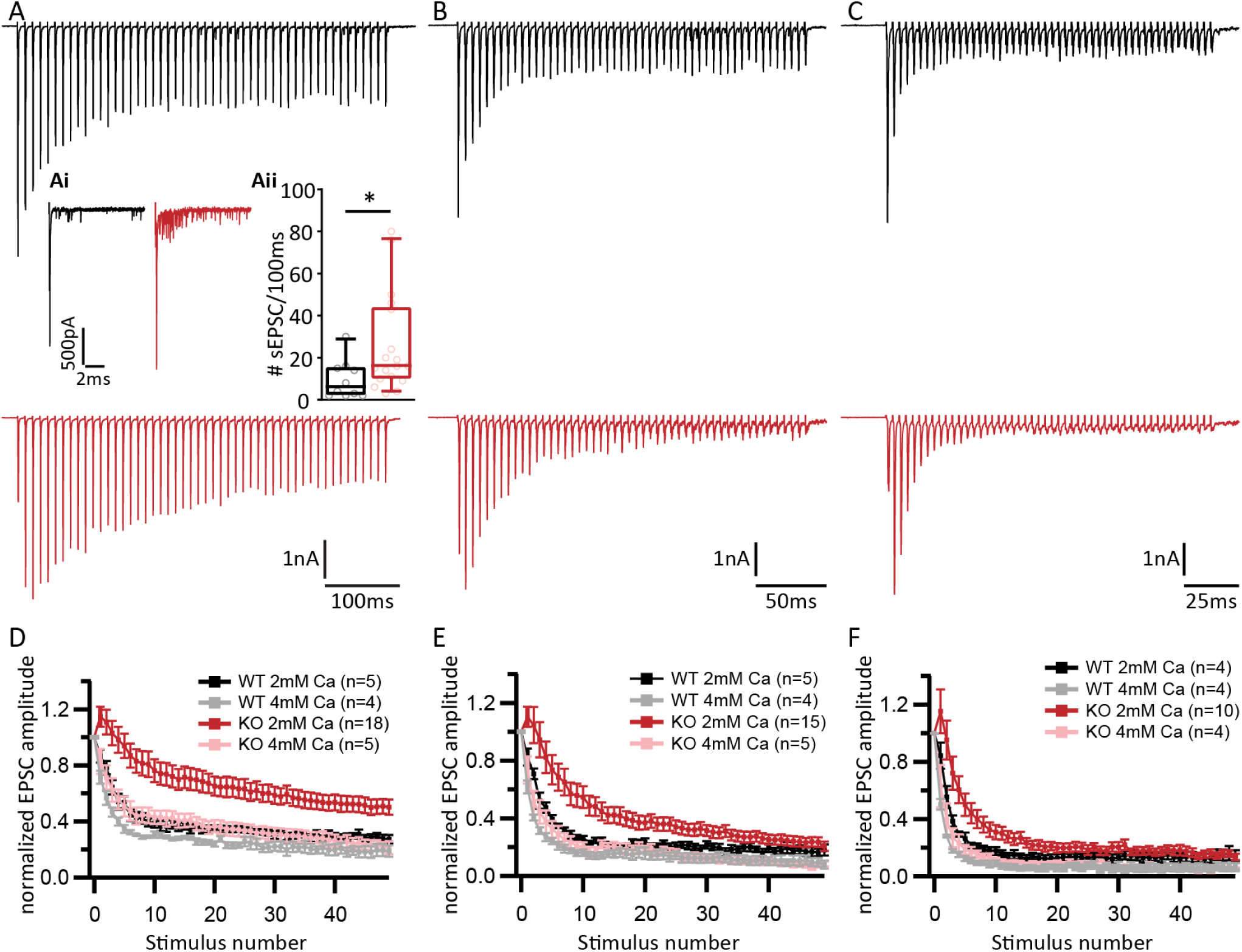
Short-term depression replaced by facilitation at endbulbs of Held in RIM-BP2 KO mice in the absence of kynurenic acid and CTZ. Representative traces of eEPSCs in response to trains of 50 action potentials delivered at frequencies of 100 (**A**), 200 (**B**) and 333Hz (**C**) recorded from WT (top, black traces) and RIM-BP2 KO (bottom, red traces). *Inset* (**Ai, Aii**): Asynchronous release calculated as the number of spontaneous EPSC events (sEPSC) within 100 ms following the synchronous release elicited by a train of 50 pulses delivered at 100Hz. A significantly higher asynchronous release was observed in the RIM-BP2 KO (red) as compared to the WT (black). Note the characteristic fast short-term depression of WT bushy cell EPSCs that is altered in the mutant. The mutant BCs show a delayed short-term depression with the first EPSC amplitude not being the largest in the train, indicating that the naive mutant synapse releases most of its vesicles later in the train. This effect is more pronounced in the higher frequencies of stimulation, as is demonstrated when the EPSC amplitudes, normalized to the first EPSC of the train are plotted against the stimulus number (**D, E, F**). This effect was abolished when extracellular Ca^2+^ was increased to 4mM (WT traces in gray and KO traces in pink).

**Figure 3.**
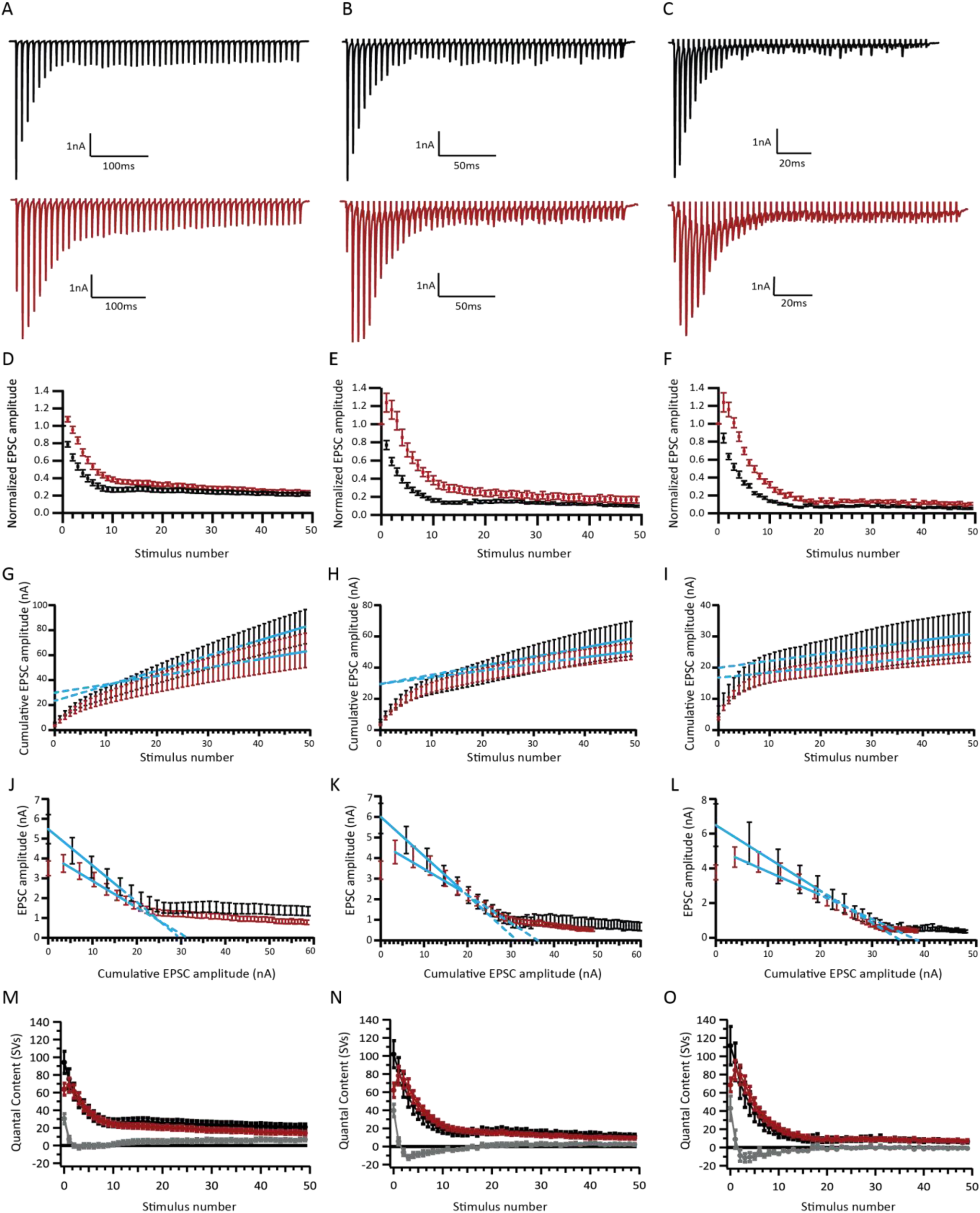
Analysis of release probability (P_r_) as well as the size and dynamics of the RRP (A-C) Representative traces of eEPSCs in response to trains of 50 action potentials delivered at frequencies of 100 (**A**), 200 (**B**) and 333Hz (**C**) recorded from WT (top, black traces) and RIM-BP2 KO (bottom, red traces). Note the characteristic fast short-term depression of WT bushy cell EPSCs that is altered in the mutant. The mutant BCs show a delayed short-term depression with the first EPSC amplitude not being the largest in the train, indicating that the naive mutant synapse releases most of its vesicles later in the train. (**D, E, F**) Average EPSC amplitudes, normalized to the first EPSC of the train plotted against the stimulus number. **(G-I)** To estimate the size of the readily releasable pool (RRP), the rate of vesicle replenishment and the release probability (P_r_) using the Schneggenburger-Meyer-Neher (SMN) method, the EPSC amplitudes of the 100 (**G**), 200 (**H**) and 333Hz (**I**) trains were plotted cumulatively against the stimulus number. The linear fit (solid blue line) to the last ten steady-state values was extrapolated to the y-axis (dotted blue line). The y-intercept value, divided by the average sEPSC amplitude yields the number of vesicles in the RRP. To calculate P_r_, the vesicle content of eEPSC1 is divided by the size of the RRP. The slope of the linear fit approximates the rate of vesicle replenishment during the train. Quantitative analysis is further elaborated in Table 1. **(J-L)** To estimate the RRP size and P_r_ using the Elmqvist and Quastel (EQ) method, absolute EPSC amplitudes were plotted against the cumulative amplitude of all the EPSCs preceding the corresponding EPSC. The linear fit to the first 3-5 points for the 100 (**J**), 200 (**K**) and 333Hz (**L**) trains (solid blue line) was forward extrapolated (dotted blue line) to the x-axis. Dividing the x-axis intercept value by the average sEPSC size, yields the size of the RRP, while the slope of the linear fit defines the P_r_. To assess whether the reduced P_r_ results from fewer “super-primed” SVs in the RRP, a subtraction analysis was performed. We first divided the average eEPSC amplitudes by the average sEPSC amplitude to calculate the SV number released during each eEPSC (quantal content) of the averaged responses to 100, 200 and 333 Hz train stimulation. Traces plot the quantal content released during each one of 50 eEPSCs in 100 (**M**), 200 (**N**) and 333Hz (**O**) trains against the stimulus number. After subtracting the quantal content for each RIM-BP2 KO eEPSC from the respective WT eEPSC, we plot the subtraction curves (gray curves at the bottom of each panel). For 100 Hz: RIM-BP2 WT N = 14; n = 16, RIM-BP2 KO N = 17; n = 21. For 200 Hz: RIM-BP2 WT N = 13; n = 16, RIM-BP2 KO N = 13; n = 16. For 333 Hz: RIM-BP2 WT N = 6; n = 12, RIM-BP2 KO N = 14; n = 17. N, number of animals; n, number of BCs.

**Table 2.** Analysis of release probability (P_r_) as well as the size and dynamics of the RRP Tau (τ) : time constant of single exponential fit to the decay of eEPSC amplitudes during the stimulation train. **eEPSC_30-50_/EPSCmax**: average amplitude of the last 20 EPSCs (30-50) in the train, normalized to the amplitude of the largest EPSC amplitude of the train. **Replenishment**: rate of vesicle replenishment. **RRP**: readily releasable pool. **P_r_**: release probability. **PPR**: paired pulse ratio, amplitude of the second EPSC of the train normalized to the amplitude of the first EPSC. Data are presented as mean ± SEM and medians are shown in parentheses. Normality of data distribution was tested with Jarque-Bera test and the variances were compared with F-test. Statistical significance of differences was assessed with unpaired Student’s t-test (shown in the table as T), when the data satisfied the criteria of normality and variance comparability. When the data did not satisfy these criteria, Mann-Whitney U-test was used instead (shown in the table as M). *p-value* < 0.05, set as threshold for statistical significance shown in bold.

| Stimulation Frequency | Parameter | RBP2 WT | RBP2 KO | p value |
| --- | --- | --- | --- | --- |
| 100Hz | eEPSC1 amplitude (nA) | 5.48 ± 0.73 (4.7) | 3.5 ± 0.37 (3.6) | **0.022, M** |
|  | tau (ms) | 33.60 ± 3.64 (27.2) | 56.95 ± 3.76 (58.03) | **0.0001, T** |
|  | eEPSC<sub>30-50</sub> (nA) | 1.29 ± 0.22 (1.09) | 0.87 ± 0.12 (0.84) | 0.1015, M |
|  | eEPSC<sub>30-50</sub>/EPSCmax | 0.23 ± 0.02 (0.21) | 0.24 ± 0.03 (0.24) | 0.8205, T |
|  | Replenishment | 2.09 ± 0.35 (1.79) | 1.44 ± 0.20 (1.2) | 0.1015, T |
|  | RRP<sub>SMN</sub> (SVs) | 398.60 ± 72.50 (341.56) | 455.61 ± 63.00 (446.32) | 0.4038, M |
|  | RRP<sub>EQ</sub>(SVs) | 575.11 ± 102.11 (504.25) | 593.62 ± 73.45 (610.12) | 0.6826, M |
| | $P_{r\text{ SMN}}$ | 0.26 ± 0.02 (0.28) | 0.17 ± 0.01 (0.15) | **0.0004, T** |
| | $P_{r\text{ EQ}}$ | 0.2 ± 0.02 (0.21) | 0.14 ± 0.01 (0.13) | **0.0336, T** |
|  | PPR | 0.79 ± 0.03 (0.72) | 1.08 ± 0.10 (0.97) | **0.0004, M** |
| 200Hz | tau (ms) | 33.05 ± 3.04 (32.46) | 70.51 ± 5.85 (65.43) | **<0.0001, M** |
|  | eEPSC<sub>30-50</sub> (nA) | 0.76 ± 0.13 (0.70) | 0.59 ± 0.07 (0.56) | 0.0608, M |
|  | eEPSC<sub>30-50</sub>/EPSCmax | 0.13 ± 0.01 (0.14) | 0.17 ± 0.03 (0.13) | 0.0608, M |
|  | Replenishment | 2.16 ± 0.37 (1.98) | 1.53 ± 0.24 (1.45) | 0.1756, T |
|  | RRP<sub>SMN</sub> (SVs) | 479.11 ± 110.23 (351.27) | 544.42 ± 77.85 (545.76) | 0.3168, M |
|  | RRP<sub>EQ</sub>(SVs) | 554.34 ± 123.03 (441.21) | 771.93 ± 108 (663.17) | 0.0883, M |
|  | P<sub>r</sub> SMN | 0.25 ± 0.02 (0.26) | 0.14 ± 0.01 (0.13) | **<0.0001, M** |
|  | P<sub>r</sub> EQ | 0.22 ± 0.02 (0.20) | 0.13 ± 0.01 (0.13) | **0.0002, T** |
|  | PPR | 0.81 ± 0.05 (0.79) | 1.28 ± 0.10 (1.21) | **<0.0001, M** |
| **333Hz** | tau (ms) | 19.34 ± 2.50 (19.77) | 39.97 ± 3.92 (40.97) | **0.0002, T** |
|  | eEPSC<sub>30-50</sub> (nA) | 0.45 ± 0.09 (0.38) | 0.48 ± 0.08 (0.41) | 0.1523, M |
|  | eEPSC<sub>30-50</sub>/EPSC<sub>max</sub> | 0.07 ± 0.01 (0.07) | 0.10 ± 0.02 (0.08) | 0.6106, M |
|  | Replenishment | 1.24 ± 0.21 (1.22) | 1 ± 0.25 (0.89) | 0.1656, M |
|  | RRP<sub>SMN</sub> (SVs) | 345.21 ± 96.42 (253.09) | 306.16 ± 36.91 (331.58) | 0.6471, M |
|  | RRP<sub>EQ</sub>(SVs) | 595.52 ± 155.74 (450.19) | 674.17 ± 76.94 (787.59) | 0.2635, M |
|  | P<sub>r</sub> SMN | 0.44 ± 0.06 (0.39) | 0.25 ± 0.03 (0.22) | **0.0043, T** |
|  | P<sub>r</sub> EQ | 0.23 ± 0.03 (0.21) | 0.15 ± 0.01 (0.15) | **0.0022, T** |
|  | PPR | 0.8 ± 0.05 (0.84) | 1.28 ± 0.11 (1.21) | **0.0009, T** |

In order to further scrutinize RRP dynamics, we studied the recovery from short-term depression, by measuring eEPSC amplitudes elicited by single stimuli presented at varying time intervals after a conditioning 100 Hz train of 50 pulses (Fig. 4). Recovery is displayed as the eEPSC amplitudes normalized to the amplitude of the first eEPSC of the conditioning train (Fig. 4A-B): RIM-BP2 deficient endbulb synapses showed a major reduction of fast recovery. The time course was fitted with a double exponential function revealing the following amplitude and tau values for the fast and slow components of recovery: A_1_ = 0.17 ± 0.05, τ_1_ = 123 ± 97.6 ms; A_2_ = 0.69 ± 0.06, τ_2_ = 6.1 ± 1.7 s for RIM-BP2 KO and A_1_ = 0.31 ± 0.04, τ_1_ = 96 ± 30 ms; A_2_ = 0.4 ± 0.04, τ_2_ = 4.6 ± 1.4 s for WT.

Next, we evaluated the impact of the impaired synaptic transmission on processing of auditory information using juxtacelluar recordings from putative spherical bushy cells (for simplicity refered to as “BCs”) *in vivo* (Fig. 5). Glass microelectrodes were stereotactically navigated to the aVCN from an occipital craniotomy, sound stimuli were presented in the open field, and BCs identified based on electrode position, first spike latency, regularity of firing and shape of the peristimulus spike time histogram (see Materials and Methods). The (non-significant) trend toward lower spontaneous firing rate (Fig. 5A, Table 3) is consistent with the reduced spontaneous ANF input (Krinner et al., 2017) and the trend toward a lower frequency of sEPSCs in RIM-BP2 KO BCs (Fig. 1-2). Sound threshold and frequency tuning were unaltered, whereby the RIM-BP2 KO data set contained more BCs with higher characteristic frequency (Fig. 5B-C, Table 3).

**Figure 5.**
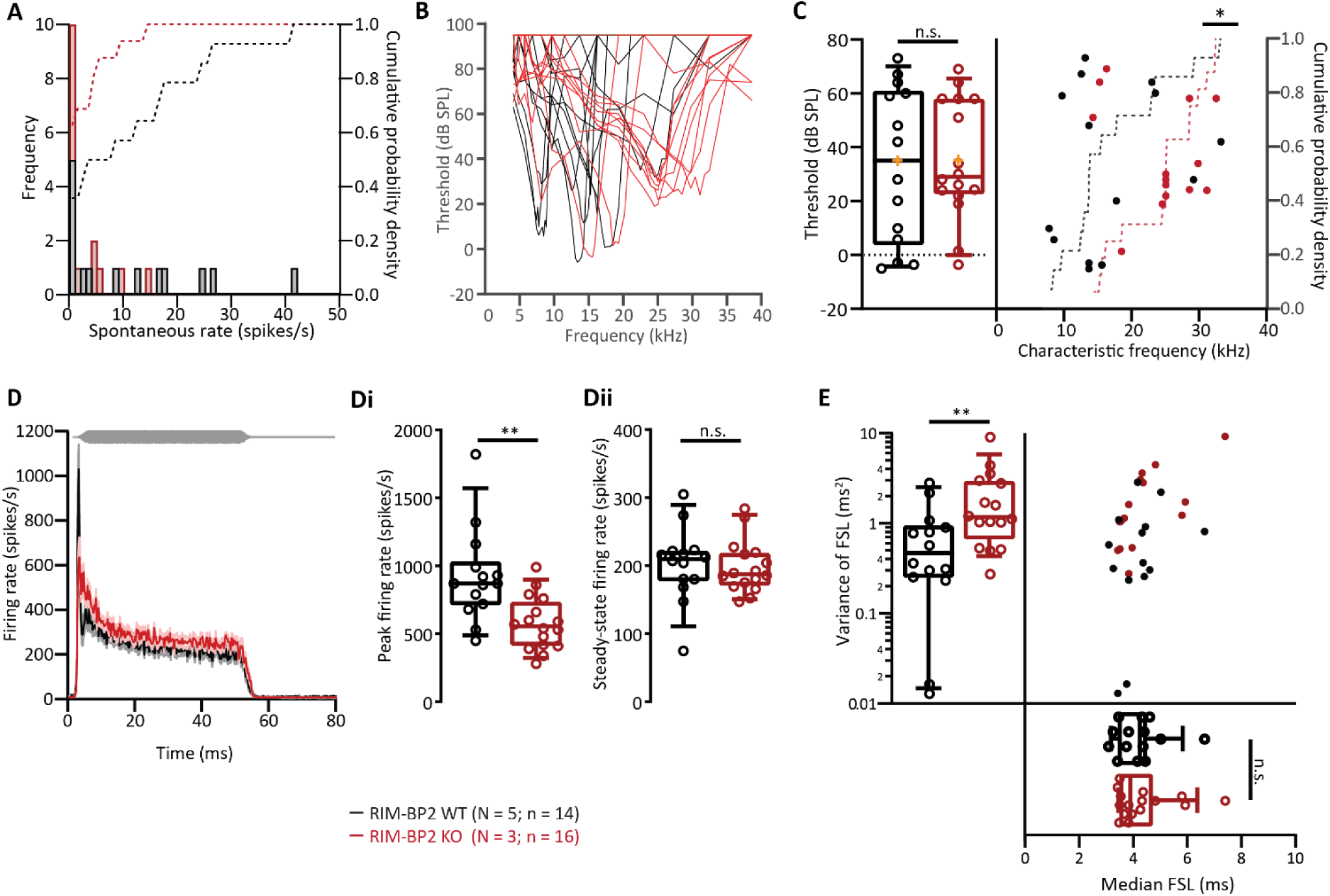
Impaired transmission of sound information in aVCN of RIM-BP2 KO mice *in vivo*. (**A**) Comparable distribution of spontaneous firing rates of single BCs in RIM-BP2 WT (black) and RIM-BP2 KO (red). The histogram represents the distribution of their frequency (left y-axis) and the dotted lines represent the cumulative probability density (right y-axis) of spontaneous firing rates. (**B-C**) Representative tuning curves of BC from RIM-BP2 WT (black) and RIM-BP2 KO (red) demonstrate preserved sharp frequency tuning and low thresholds at the characteristic frequencies (frequency for which spike rate increase requires least sound intensity, **C,** *left*) in RIM-BP2 KO BCs. For unknown reasons, we encountered more BCs with high characteristic frequency in RIM-BP2 KO (**C,** *right*). (**D**) Rise-aligned peristimulus time histogram (PSTH) of the BC response to 50 ms tone burst stimulation (at characteristic/best frequency, 30 dB above threshold, stimulus represented in gray) in RIM-BP2 WT (black; N=5; n=14) and RIM-BP2 KO (red; N=3; n=16). PSTH presented as mean (solid lines) ± SEM (shaded area). Peak onset firing rate was significantly reduced in RIM-BP2 KO BCs (**Di**) while the steady-state firing rate was comparable between the two genotypes (**Dii**). Variance in the first spike latency of PSTH (in D) was increased in RIM-BP2 KO units while the median first spike latency remained unperturbed (**E**). Data information: Significance levels: n.s. p ≥ 0.05, *p < 0.05, **p < 0.01; n = number of BCs and N = number of mice. Box and whisker plot represents median, lower/upper quartiles and 10-90^th^ percentiles. Each data point represents the response of a BC. For details about mean ± SEM, median, sample size and statistics see Table 3.

**Table 3.** Analysis of *in vivo* extracellular recordings from single BCs. Data are presented as mean ± SEM and medians are shown in parentheses. Normality of data distribution was tested with Kolmogorov-Smirnov test and the variances were compared with F-test. Statistical test used to assess the significance of differences is indicated in the column of p-value. Unpaired Student’s t-test (shown in the table as T) was used, when the data satisfied the criteria of normality and variance comparability. Normally distributed data with unequal variances were compared using Student’s t-test with Welch’s correction (shown in the table as TW). Non-normally distributed data were tested with Mann-Whitney U-test or Kolmogorov-Smirnov test (shown in the table as M and K respectively). *p-value* < 0.05, set as threshold for statistical significance shown in bold.

| Parameter | RIM-BP2 WT<br>(N = 5; n = 14) | RIM-BP2 KO<br>(N = 3; n = 16) | p value |
| --- | --- | --- | --- |
| Spontaneous rate (spikes/s) | 10.90 ± 3.42 (5.88) | 2.74 ± 1.07 (0.60) | 0.18, M |
| Threshold (dB SPL) | 33.20 ± 7.76 (35.00) | 35.20 ± 5.50 (29.00) | 0.83, T |
| Characteristic frequency (kHz) | 16.75 ± 2.06 (13.66) | 24.21 ± 1.59 (25.05) | **0.015, K** |
| Peak firing rate (spikes/s) | 922.10 ± 91.65 (870.00) | 575.60 ± 49.24 (555.00) | **0.0033, TW** |
| Steady-state firing rate (spikes/s) | 202.30 ± 14.53 (209.80) | 197.80 ± 9.6 (187.40) | 0.79, T |
| Variance of FSL (ms<sup>2</sup>) | 0.76 ± 0.22 (0.47) | 2.11 ± 0.56 (1.18) | **0.008, M** |
| Median FSL (ms) | 4.20 ± 0.24 (4.25) | 4.36 ± 0.28 (3.90) | 0.93, M |

The peak firing rate at sound onset was significantly reduced in RIM-BP2 KO BCs (Fig. 5D, by approximately 40%, Table 3) compatible with the reduced initial release probability. The adapted firing rates were not significantly reduced (Fig. 5Dii, Table 3), which is consistent the better maintained EPSC amplitudes during steady state response to train stimulation in BCs (RIM-BP2 WT: 1.2 ± 0.2 nA, RIM-BP2 KO: 0.9 ± 0.1) (Fig. 3A-F, Table 2) and the normal adapted firing rate of ANFs (Krinner et al., 2017).

In addition, we found the temporal jitter of the first spike after stimulus onset to be greater in RIM-BP2 KO BCs, while the first spike latency was comparable between the two genotypes (Fig. 5E, Table 3). The stronger reduction of the peak rate and increased first spike latency jitter of BCs in comparison to ANF likely reflects the impaired transmission at the RIM-BP2-deficient endbulb, which degrades information processing in the lower auditory pathway beyond what is caused by the mildly affected synaptic sound encoding in the cochlea. This hypothesis is further supported by the auditory brainstem responses of RIM-BP2 KO mice, which show a more pronounced amplitude decline for the aVCN related wave III than for the auditory nerve related wave I (Krinner et al., 2017)). In conclusion, the reduced release probability of the endbulbs impairs the transmission of sound onset information, which is likely to hamper hearing and auditory tasks such as gap detection and sound localization in particular.

### RIM-BP2 disruption appears not to alter the molecular composition of the endbulb active zones

Given the scope of protein interactions of RIM-BP that includes Ca^2+^ channels, large conductance Ca^2+^ activated K^+^ channels (Sclip et al., 2018) and multi-domain proteins of the AZ, we considered the possibility that some of the above described physiological alterations might reflect changes in the abundance of other AZ proteins. In order to test for potential effects of RIM-BP2 disruption on the molecular composition of the AZs, we performed semi-quantitative, confocal immunofluorescence microscopy in coronal brain slices. RIM-BP2 WT and RIM-BP2 KO samples were harvested and processed strictly in parallel. Likewise, images were acquired using the same laser power and gain settings at the same confocal microscope. Excitatory AZs facing the postsynaptic BC were identified by co-localization of immunofluorescence of the targeted AZ protein with the immunofluorescence of the vesicular glutamate transporter VGlut1 or a juxtaposition to the immunofluorescence of homer 1, a scaffold of excitatory synapses and a lack of juxtaposition to immunofluorescence of gephyrin, a scaffold of inhibitory synapses (Fig. 6). We focused our analysis on the spherical or ovoid BC soma which are engaged by a corona of synapses. Staining for RIM-BP2 showed the expected corona of immunofluorescence spots in WT slices, but no obvious synaptic immunofluorescence in RIM-BP2 KO slices (Fig. 6A, Fig. 6B, left: integrated fluorescence within the Vglut1 positive volume, right: lack of spots). We did not observe significant differences in the integrated immunofluorescence or the number of puncta of Munc13-1 (Fig. 6C, D), RIM2 (Fig. 6E, F), CAST (Fig. 6G, H), and bassoon (Fig. 6I, J) in RIM-BP2 KO slices, suggesting an unaltered abundance of these multi-domain proteins at the excitatory AZ facing the BC in the absence of RIM-BP2.

**Figure 6.**
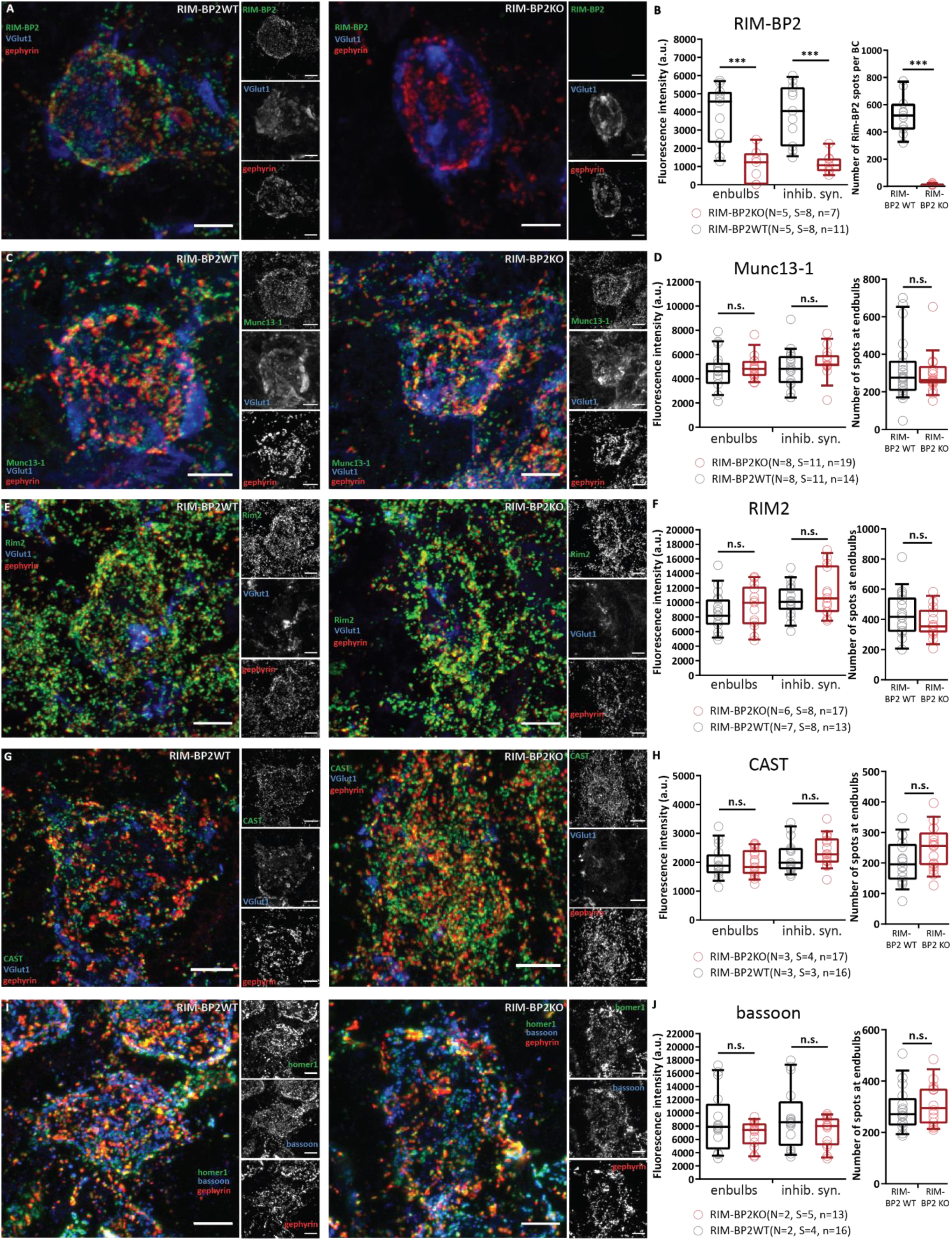
RIM-BP2 disruption does not seem to alter the molecular composition of the endbulb active zones. (**A, C, E, G, I**) Representative maximal z-projections of confocal image stacks of BCs in RIM-BP2 WT shown on left and RIM-BP2 KO on right. 30 μm coronal brainstem slices were immunolabelled for RIM-BP2 (**A**), Munc13-1 (**C**), RIM2 (**E**) and CAST (**G**) and co-stained for VGlut1 to outline endbulbs and gephyrin for inhibitory synapse labelling. For bassoon immunolabelling (**I**) we co-stained for homer1 (excitatory postsynapses) and gephyrin. (**B, D, F, H, J**) Quantification of fluorescence intensity of CAZ proteins at endbulbs and inhibitory synapses of BCs (*left)*, and the number of spots of the targets CAZ protein at endbulbs (*right*). (**A**) Representative z-projection shows loss of RIM-BP2 immunoreactivity in the RIM-BP2 KO brainstem sliced. There was only a faint residual and likely unspecific signal remaining at excitatory and inhibitory synapses (**B**, left), RIM-BP2 immunofluorescent spots were nearly abolished (**B**, right). The immunofluorescence intensity of Munc13-1 (**D**), RIM2 (**F**), CAST (**H**) and bassoon (**J**) were unaltered at RIM-BP2 KO endbulbs. Right columns of D, F, H, J show similar number of CAZ protein spots localized at KO endbulb AZs as in the WT AZs. The numbers were calculated by subtracting the gephyrin colocalized spots from the total number of spots per BC. The data are presented as box and whiskers plots (grand median of mean estimates for all BCs, lower/upper quartiles, 10-90^th^ percentiles). Each data point represents the mean estimate of fluorescence intensity of the active zones of each BC included in the analysis. Statistical significance of differences between groups was determined with unpaired Student’s t-test (with Welch’s correction, when variances differed significantly), if the data’s distribution did not differ from a normal distribution or with Mann-Whitney U test in case of non-normally distributed data. Normality of distribution was tested with Jarque-Bera test and variances were compared with F-test. n.s. p-value ≥ 0.05. Samples from RIM-BP2 WT and RIM-BP2 KO mice, aged p15-p21 were harvested and processed strictly in parallel and images were acquired in parallel using the same laser power and gain settings at the same confocal microscope. Data information: *N*: number of animals, *S*: number of slices, *n*: number of BCs. All scale bars: 5μm

### Increased Ca^2+^ influx improves release probability but unmasks impaired SV replenishment during train stimulation

Unaltered Ca^2+^ influx and RRP size but reduced P_r_ led us to focus the analysis on the coupling of Ca^2+^ channels to SV release. As a first approach we increased the presynaptic Ca^2+^ influx by elevating the extracellular Ca^2+^ concentration [Ca^2+^]_e_ from physiological (2mM) to 4mM. This manipulation abolished the differences in P_r_ (both time course of depression and PPR were WT-like, Fig. 7, Table 4). This is consistent with a greater diffusional distance between Ca^2+^ channels and SV release sites that can be overcome when more Ca^2+^ enters per channel opening. Alternatively, or in addition, greater Ca^2+^ influx might foster Ca^2+^ dependent priming or facilitation of release. In addition, increased Ca^2+^ influx unmasked a slowed SV replenishment during train stimulation in RIM-BP2 deficient endbulbs.

**Figure 7.**
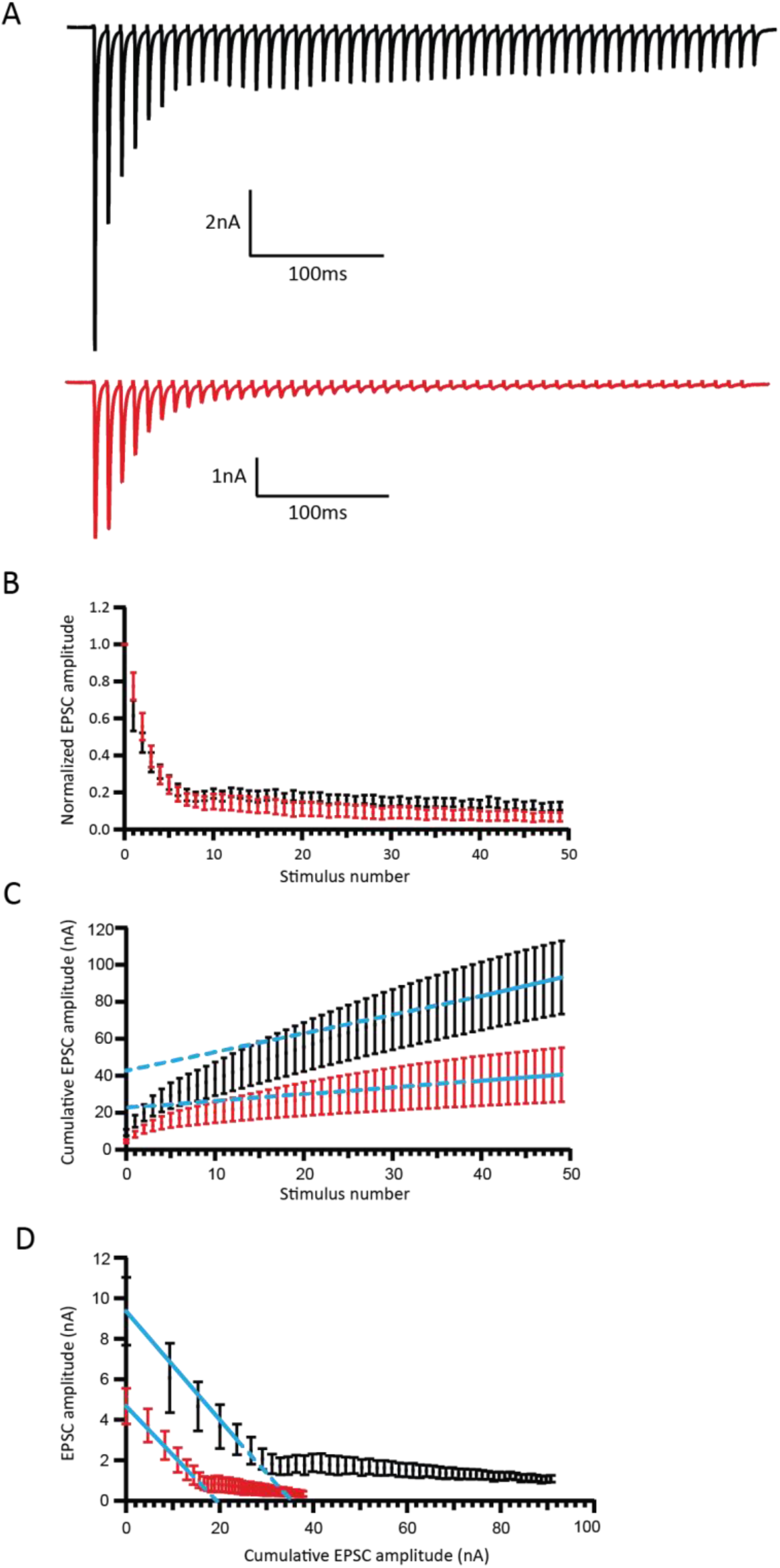
Analysis of release probability (P_r_) as well as the size and dynamics of the RRP at 4 mM [Ca^2+^]_e_. (**A**) Representative traces of eEPSCs in response to trains of 50 action potentials delivered at a 100 Hz frequency, recorded from WT (top, black traces) and RIM-BP2 KO (bottom, red traces). When the mutant terminals are exposed to 4mM [Ca^2+^]_e_, a WT-like depression pattern is restored. (**B**) This is even more obvious when the mean EPSC amplitudes, normalized to the first EPSC of the train are plotted against the stimulus number. (**C**) We estimate the size of the readily releasable pool (RRP), the rate of vesicle replenishment during the train and the release probability (P_r_) using the SMN method. (**D**) We estimate the RRP size and P_r_ using the Elmqvist and Quastel (EQ) method. For 100 Hz: RIM-BP2 WT N = 7; n = 7, RIM-BP2 KO N = 5; n = 5. N, number of animals; n, number of BCs. Quantitative analysis is further elaborated in Table 4.

**Table 4.**
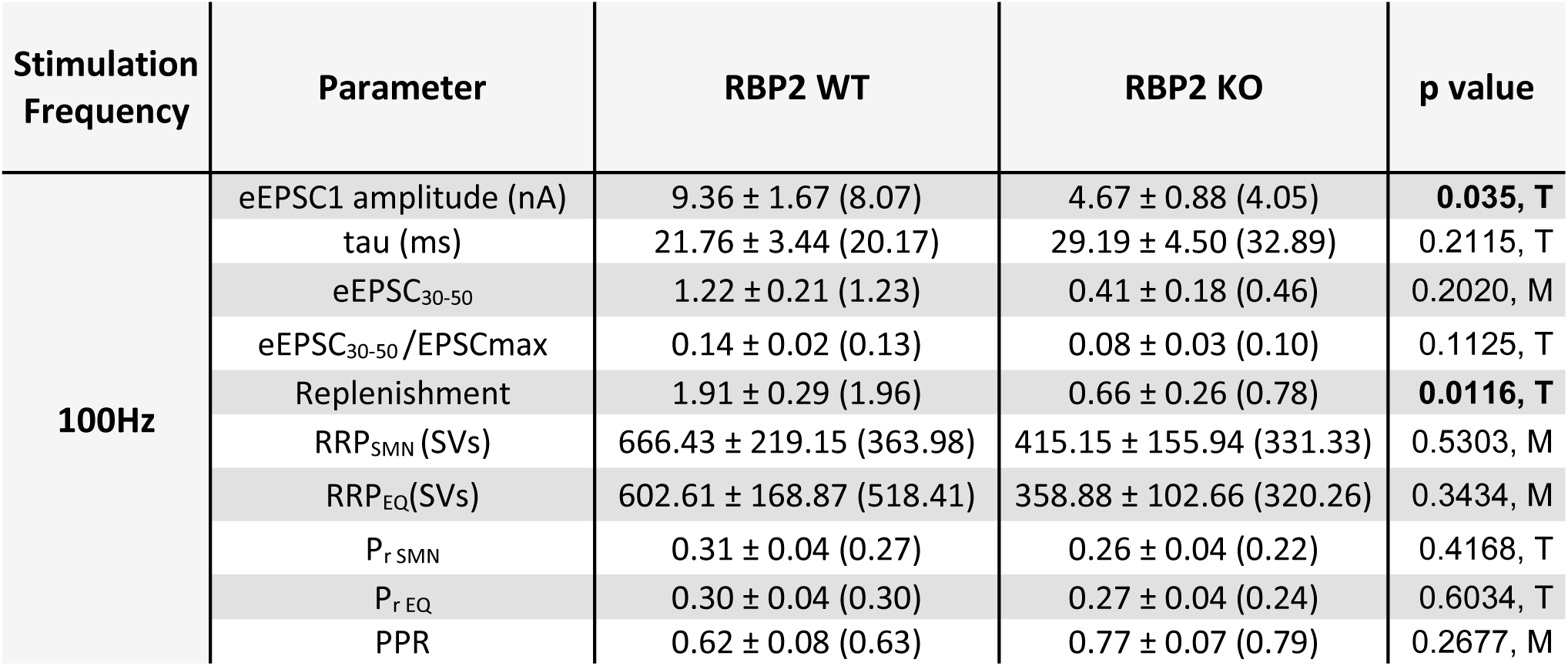
Analysis of release probability (P_r_) as well as the size and dynamics of the RRP at 4 mM [Ca^2+^]_e_ Tau (τ) : time constant of single exponential fit to the decay of eEPSC amplitudes during the stimulation train. **eEPSC_30-50_/EPSCmax**: average amplitude of the last 20 EPSCs (30-50) in the train, normalized to the amplitude of the largest EPSC of the train. **Replenishment**: rate of vesicle replenishment during the train. **RRP**: readily releasable pool. **P_r_**: release probability. **PPR**: paired pulse ratio, amplitude of the second EPSC of the train normalized to the amplitude of the first EPSC. Data are presented as mean ± SEM and medians are shown in parentheses. Normality of data distribution was tested with Jarque-Bera test and the variances were compared with F-test. Statistical significance of differences was assessed with unpaired Student’s t-test (shown in the table as T), when the data satisfied the criteria of normality and variance comparability. When the data did not satisfy these criteria, the Mann-Whitney U-test was used instead (shown in the table as M). *p-value* < 0.05, set as threshold for statistical significance shown in bold.

### RIM-BP2 disruption alters the topography of Ca^2+^ channels at endbulb active zones

In order to probe for potential morphological correlates of altered release probability we employed immunofluorescence microscopy and electron microscopy of immunolabeled Ca^2+^ channels. First, we performed a strictly parallel study of Ca_V_2.1 distribution at excitatory synapses around the BCs of RIM-BP2 KO and WT mice using stimulated emission depletion (STED) nanoscopy (Fig. 8A-K) as well as confocal microscopy (as described above, Fig. 8-1A, B). We used bassoon and homer1 as context markers to analyse the presynaptic Ca_V_2.1 immunofluorescence distribution. Both the integrated Ca_V_2.1 immunofluorescence intensity per AZ and the number of Ca_V_2.1 puncta were unaltered at the confocal level (Fig. 8-1B), which is consistent with finding normal presynaptic Ca^2+^ influx (Fig. 2). For our 2-colour STED analysis we focused on AZ/PSD appositions (Ca_V_2.1/homer1, bassoon/homer1) of endbulbs of Held.

**Figure 8.**
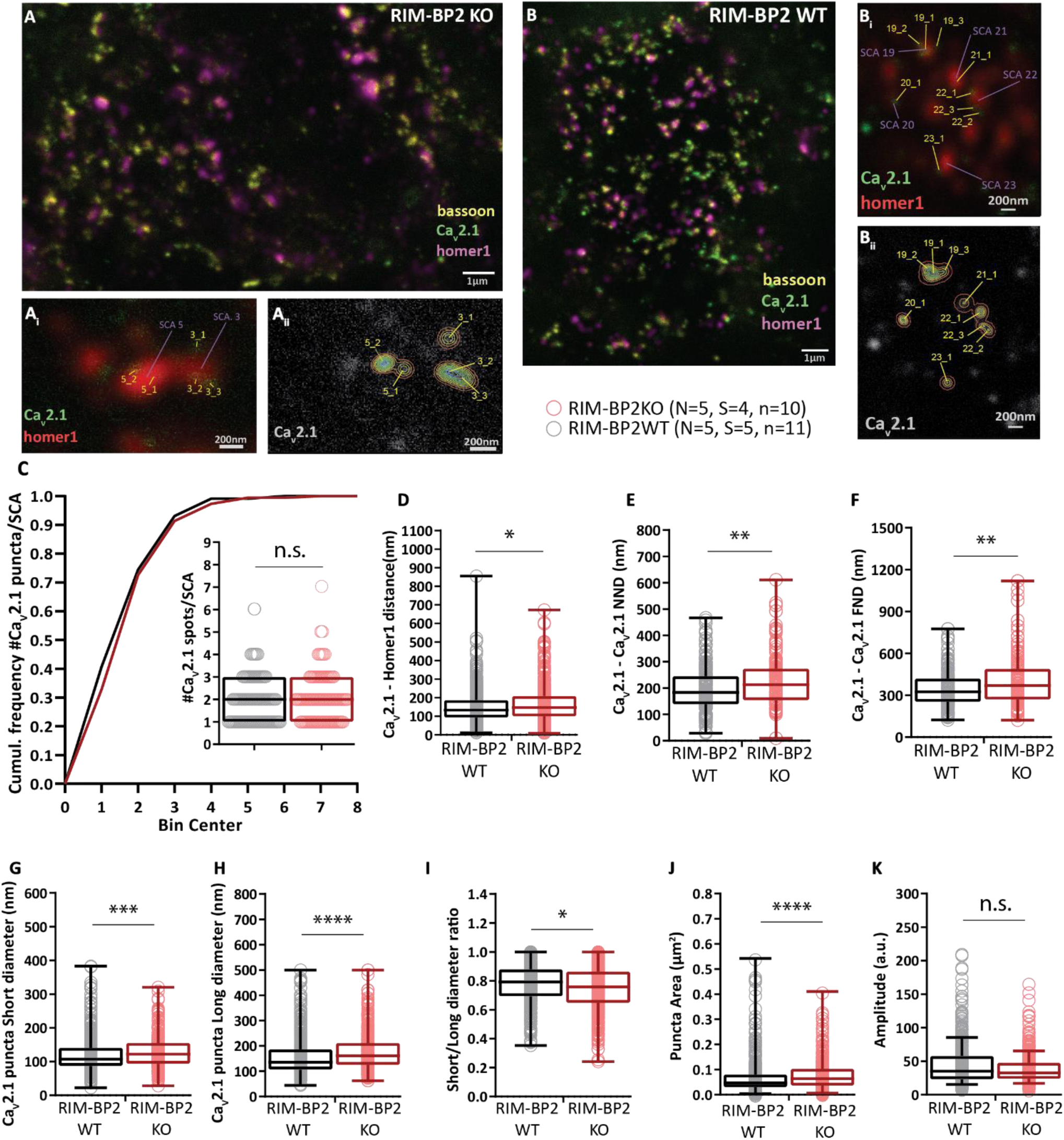
RIM-BP2 disruption alters the topography of Ca^2+^ channels at endbulb active zones. (**A-B**) Analysing sections from 20 µm thick brain slices stained for Ca_V_2.1 (STED), bassoon (STED) and homer1 (Confocal) with STED nanoscopy, uncovered differences in the topography and dimensions of Ca_V_2.1 clusters at AZ – to – PSD SCAs of RIM-BP2 WT (**A, A_i_, A_ii_**) and RIM-BP2 KO (**B, B_i_, B_ii_**) endbulb terminals. (**C**) The number of clusters per SCA is unchanged in the KO. In the absence of RIM-BP2, the Ca_V_2.1 puncta (yellow tags **A_i_, A_ii_, B_i_, B_ii_**) are located further from the center of the SCA (**D**), defined as the center of the Ηomer1 puncta (violet tags **A_i_, B_i_**). The clusters are also located further apart from each other, shown by increased Nearest Neighbour (**E**) and Furthest Neighbour (**F**) distances within SCAs. (**G, H, J**) 2D – Gaussian fitting yielded the short and long cluster diameters at half maxima. (**I**) The deletion of RIM-BP2 leads to larger and more elongated Ca_V_2.1 puncta at the endbulb of Held active zones. (**K**) The amplitude of fluorescence at the center of Ca_V_2.1 puncta does not differ significantly between RIM-BP2 WT and RIM-BP2 KO active zones. Normality was tested with Jarque-Bera test. Statistical significance between groups was tested with Mann-Whitney U-test for non-normally distributed data or with Student’s t-test. \*\*\*\**p-value* < 0.0001, \*\*\**p-value* < 0.001, \*\**p-value* < 0.01, \**p-value* < 0.05. *N*: number of animals, *S*: number of slices, *n*: number of BCs.

**Figure 8-1:**
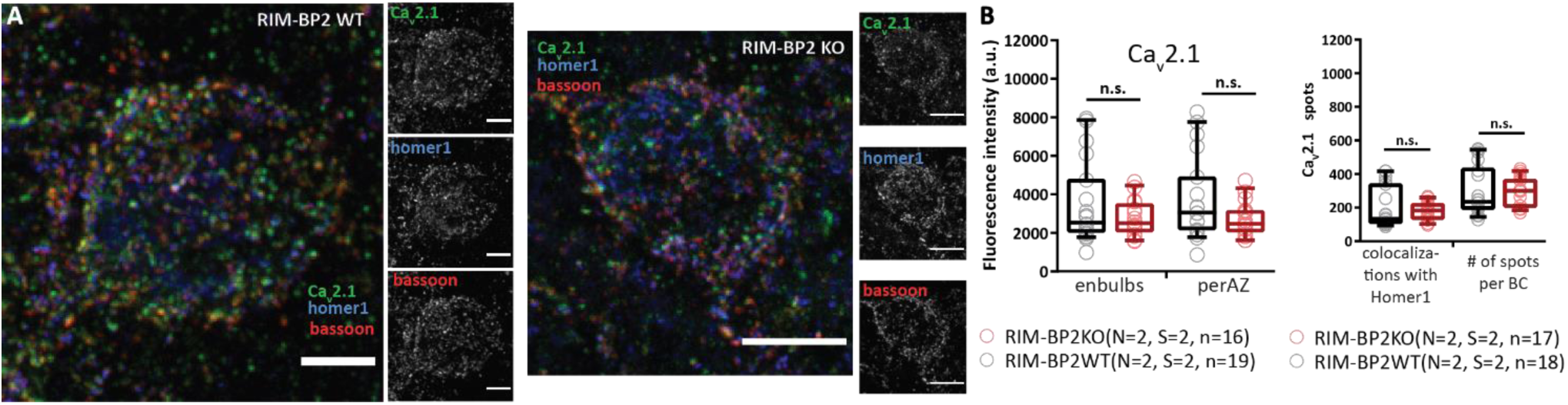
Unaltered Ca_V_2.1 immunofluorescence intensity in RIM-BP2-deficient endbulbs of Held. (**A**) Maximal z-projections from confocal image stacks display the immunolabelling against Ca_V_2.1, bassoon and homer1 at endbulb AZs in coronal brain-stem slices of mouse aVCN. (**B**) No quantitative change was found in the integrated immunofluorescence (*left*) or number (*right*) of Ca_V_2.1 puncta in either the excitatory endbulb AZs (juxtaposed to homer1 immunofluorescence) or all AZs (endbulb + inhibitory AZ) facing BC of RIM-BP2 KOs. \**p-value* < 0.05. *N*: number of animals, *S*: number of slices, *n*: number of BCs.

We interpret the Ca_V_2.1 and bassoon puncta discerned by STED to represent individual AZs. As the PSD was studied at confocal resolution, we assume the larger homer1-spots represent a merger of several small PSDs. We operationally defined the organization of several Ca_V_2.1 or bassoon puncta around a single homer1 spot a synaptic contact assembly (SCA) whereby its center corresponds to the center of the homer1 punctum. Our analysis indicated a wider spatial distribution of Ca_V_2.1 puncta in RIM-BP2 KO which localized further away from the SCA center and from each other (Fig. 8A-K), suggestive of a more dispersed Ca_V_2.1 topography. Similar findings were made for bassoon (Fig. 8-2).

**Figure 8-2.**
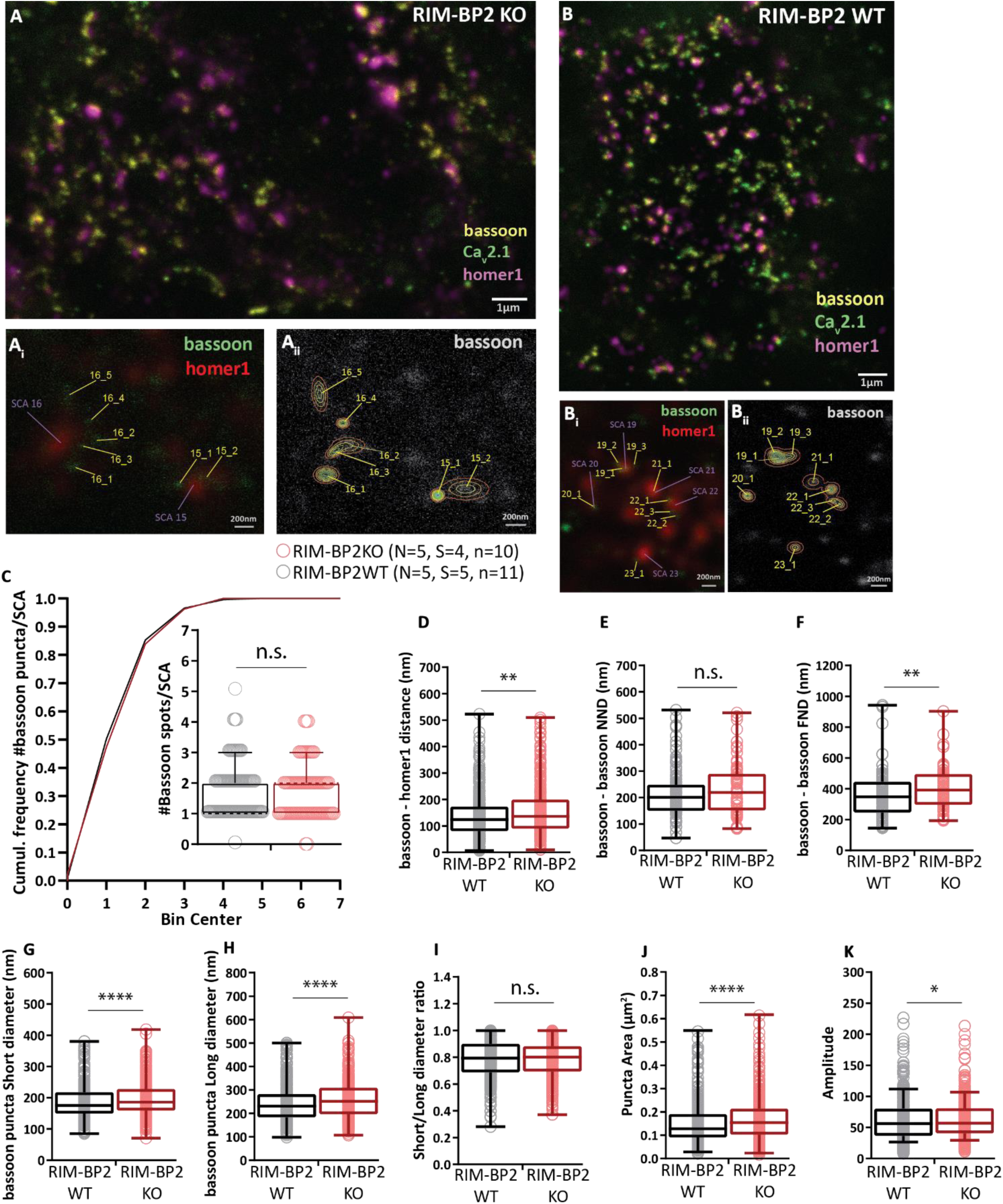
RIM-BP2 disruption alters distribution and extent of bassoon clusters marking the presynaptic density of endbulb active zones. Analysing sections (**A, B**) from 20 µm thick brain slices stained for Ca_V_2.1 (STED), bassoon (STED) and homer1 (Confocal) with super-resolution STED microscopy, uncovered differences in the amplitude of fluorescence, topography and dimensions of bassoon positive puncta in the absence of RIM-BP2. The bassoon puncta were detected and fitted with 2D – Gaussian functions at AZ – to – PSD SCAs of RIM-BP2 WT (**A, A_i_, A_ii_**) and RIM-BP2 KO (**B, B_i_, b_ii_**) endbulb terminals. The number of puncta per SCA is unchanged in the KO (**C**). In the absence of RIM-BP2, the bassoon puncta (yellow tags **A_i_, A_ii_, B_i_, B_ii_**) are located further from the center of the SCA (**D**), defined as the center of the postsynaptic homer1 puncta (violet tags **A_i_, B_i_**). We found no significant difference in the Nearest Neighbour distance (E). Furthest neighbour distance (**F**) was significantly increased in the mutant. Comparing the short and long puncta diameters at half maxima, shows that the deletion of RIM-BP2 leads to proportionally (no change in short/long diameter ratio, **I**) larger (**G, H, J**) bassoon puncta at endbulb of Held active zones. The amplitude of fluorescence at the center of bassoon puncta is significantly increased RIM-BP2 KO active zones (**K**). Normality was tested with Jarque-Bera test. Statistical significance between groups was tested with Mann-Whitney U-test for non-normally distributed data and with Student’s t-test for data showing a normal distribution. \*\*\*\**p-value* < 0.0001, \*\*\**p-value* < 0.001, \*\**p-value* < 0.01, \**p-value* < 0.05. *N*: number of animals, *S*: number of slices, *n*: number of BCs.

We then turned to immunolabelling of Ca_V_2.1 in SDS-treated freeze-fracture replica (SDS-FRIL, (Nakamura et al., 2015)). To image the endbulbs onto the BCs, we focused on the BC rich rostral-most aVCN sections of the brainstem. Endbulb terminals were prominently distinguishable in our replicas by their large size synapsing on to the BC soma (Fig. 9-1A). We also validated that we were imaging the correct area and cell type, by analysing the intramembrane particle (IMP) clusters representing PSDs on the exoplasmic face (E-face) of the BC soma (Fig. 9-1C-F). Our estimates of PSD areas (Fig. 9-1B, Table 5) were comparable to the ones previously reported for ANF-BC synapses (Rubio et al., 2017). We then assessed the protoplasmic face (P-face) of the replicas for the analysis of AZ proteins and Ca_V_2.1 channel distribution (Fig. 9-2). AZs were located by simultaneous immunolabelling of three characteristic AZ proteins: RIM, neurexin, and ELKS with 5 nm gold particles. The number of AZ particles was less than that previously observed in other types of synapses (Miki et al., 2017) which might reflect lower expression of ELKS in the endbulb synapses. Since the samples from both genotypes were handled simultaneously by the same experimenter, the comparison between the genotypes remains valid. Nonetheless, given the low labelling efficiency for AZ proteins, we used AZ markers primarily to identify the location of AZs. AZ area was characterized by IMPs of distinct shape, number and size compared with those in surrounding areas, and demarcated manually by connecting the outermost IMPs (Fig. 9-2B-C). The estimated AZ area was comparable between RIM-BP2 KO and WT (Fig. 9-2D). When analysing the distribution of Ca_V_2.1 channels labelled with 10 nm gold particles within the AZ area, both the number and density of Ca_V_2.1 particles were significantly reduced in the RIM-BP2 KO (Fig. 9-2G-H, Table 5).

**Figure 9.**
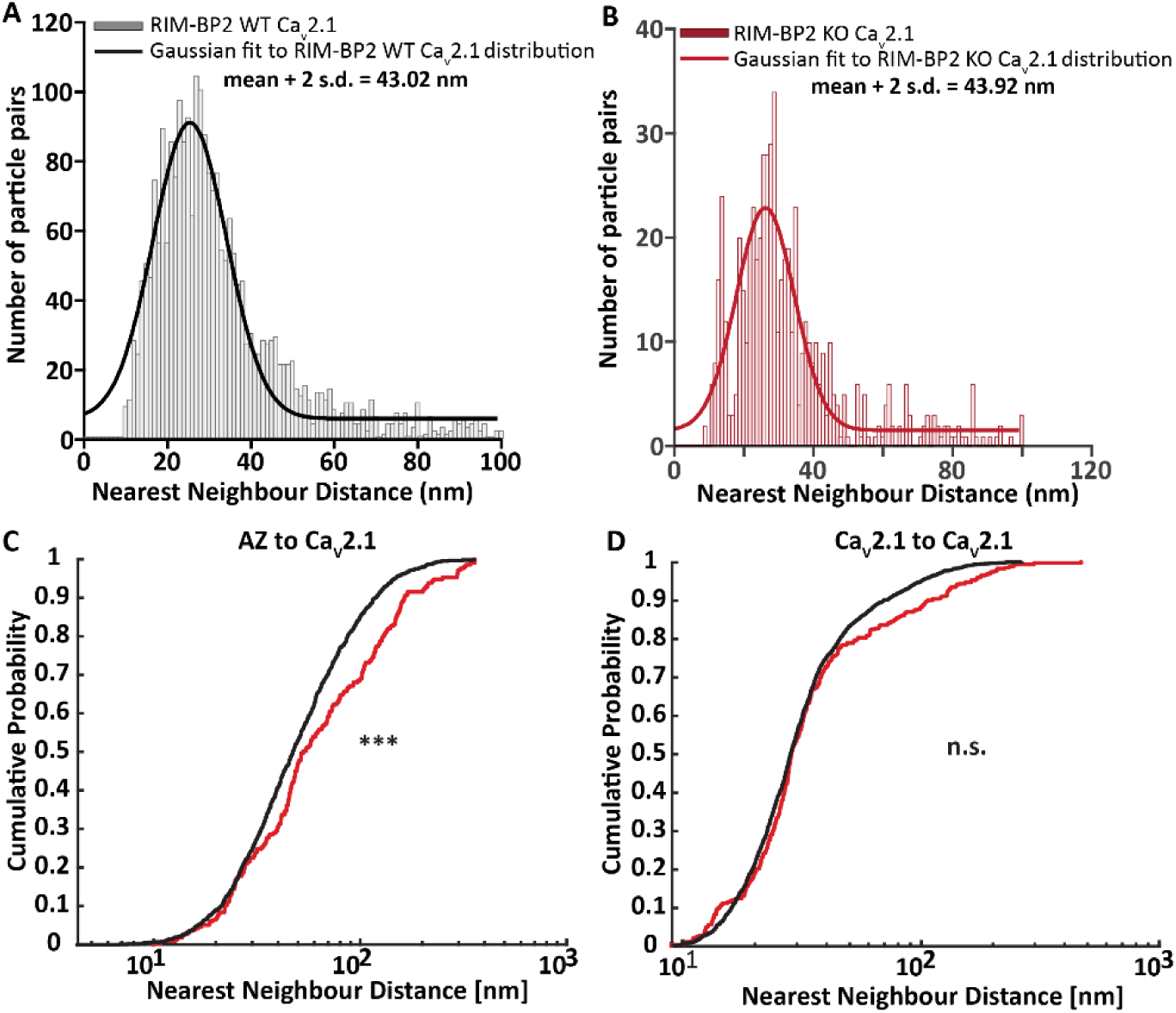
Altered topography of Ca_V_ channels relative to AZ proteins. (**A-B**) Histogram of nearest neighbor distances (NNDs) between Ca_V_2.1 to Ca_V_2.1 gold particles in RIM-BP2 WT (**A**) and KO (**B**). The solid bold line represents Gaussian curve fitted to the NND distribution in RIM-BP 2 WT (black, **A**) and KO (red, **B**). (**C-D**) Cumulative probability of NNDs between AZ to Ca_V_2.1 particles (**C**) and Ca_V_2.1 to Ca_V_2.1 particles (**D**) in RIM-BP2 WT (black) and KO (red). ns: not significant, *** *p-value* < 0.001. For details about mean ± SEM, median, sample size and statistics see Table 5.

**Figure 9-1.**
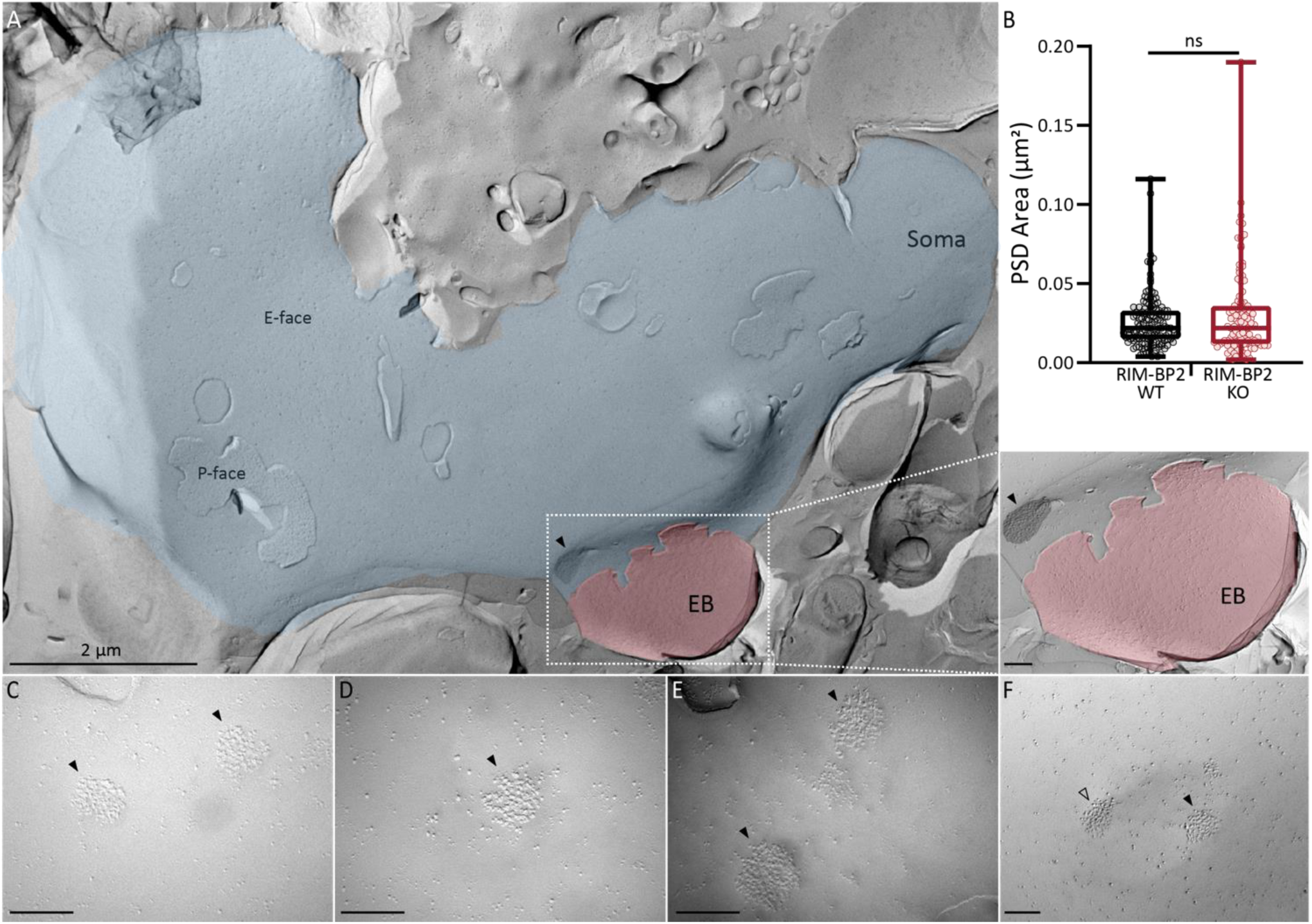
Identification of endbulb of Held synapses. (**A**) SDS-FRIL electron micrograph at low magnification (9700x) showing E-face of a bushy cell (BC) soma (blue) with patches of P-face, contacted by an endbulb (red). IMP-cluster representing the PSD of auditory nerve on the BC soma highlighted in black with a solid black arrow head. Inset shows the magnified (97,000x) view of the endbulb synapse on to the BC with a PSD IMP-cluster. (**B**) Comparable PSD areas at endbulbs of Held in RIM-BP2 WT and KO. (**C-F**) High magnification (C-E 93,000x; F 97,000x) images of IMP-clusters of BC soma facing the endbulb. PSD IMP-clusters indicated by solid black arrow heads. Open black arrow head in F marks the PSD IMP-cluster that lies on the curvature of the synaptic cleft depression and hence was not included in the analysis. All unmarked scale bars are 200 nm. For details about PSD values, sample size and statistics see Table 5.

**Figure 9-2.**
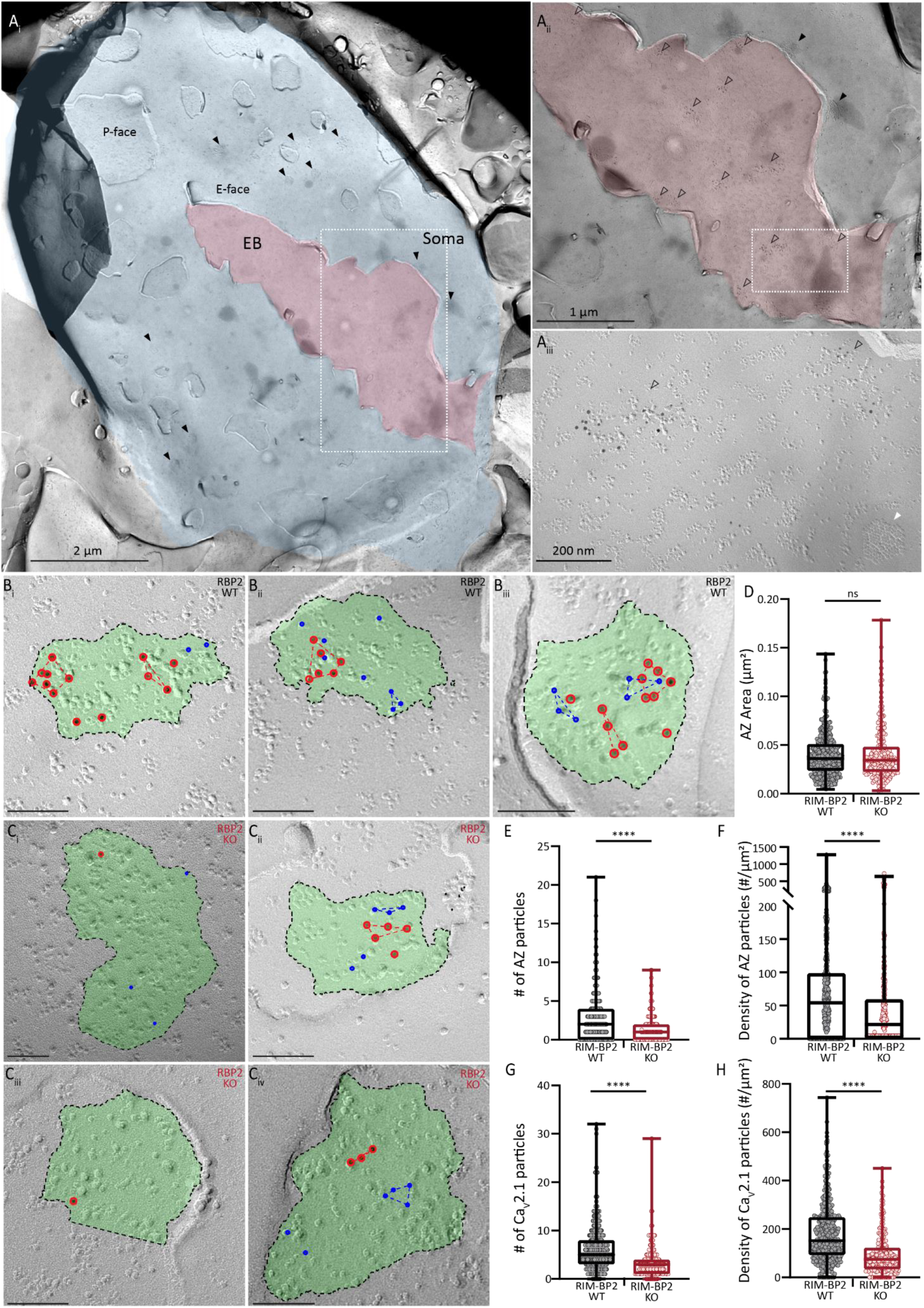
Similar AZ size but reduced number of AZ and Ca_V_2.1 labels in endbulbs of RIM-BP2 KO. (**A**) SDS-FRIL electron micrograph at low magnification (9700x) showing E-face of a bushy cell (BC) soma (blue) with patches of P-face, connected to an endbulb (red). (**Ai**) Some of the IMP-clusters representing the PSDs of ANF on the BC soma highlighted by solid black arrow heads. (**Aii**) Magnified (18,500x) view of the box in Ai of endbulb synapse (red) on to the BC soma (blue) with IMP-clusters for AZ and PSD indicated by open and solid black arrows respectively. (**Aiii**) Magnified (97,000x) view of the box in Aii with IMP-clusters for AZ indicated by open arrows and P-face of a gap junction indicated by a solid white arrow. (**B-C**) FRIL images showing endbulb AZs (green) in RIM-BP2 WT (B) and KO (C), immunolabelled for AZ molecules (RIM, neurexin, ELKS) and Ca_V_2.1 channels with 5 nm (blue circles) and 10 nm (red circles) gold particles respectively. (**D-H**) Quantitative analysis of AZ area (**D**), number of AZ (**E**) and Ca_V_2.1 (**G**) gold particles, and density of AZ (**F**) and Ca_V_2.1 (**H**) gold particles. All unlabelled scale bars are 100 nm. ns: not significant, **** *p-value* < 0.0001. For details about mean ± SEM, median, sample size and statistics see Table 5.

**Table 5.**
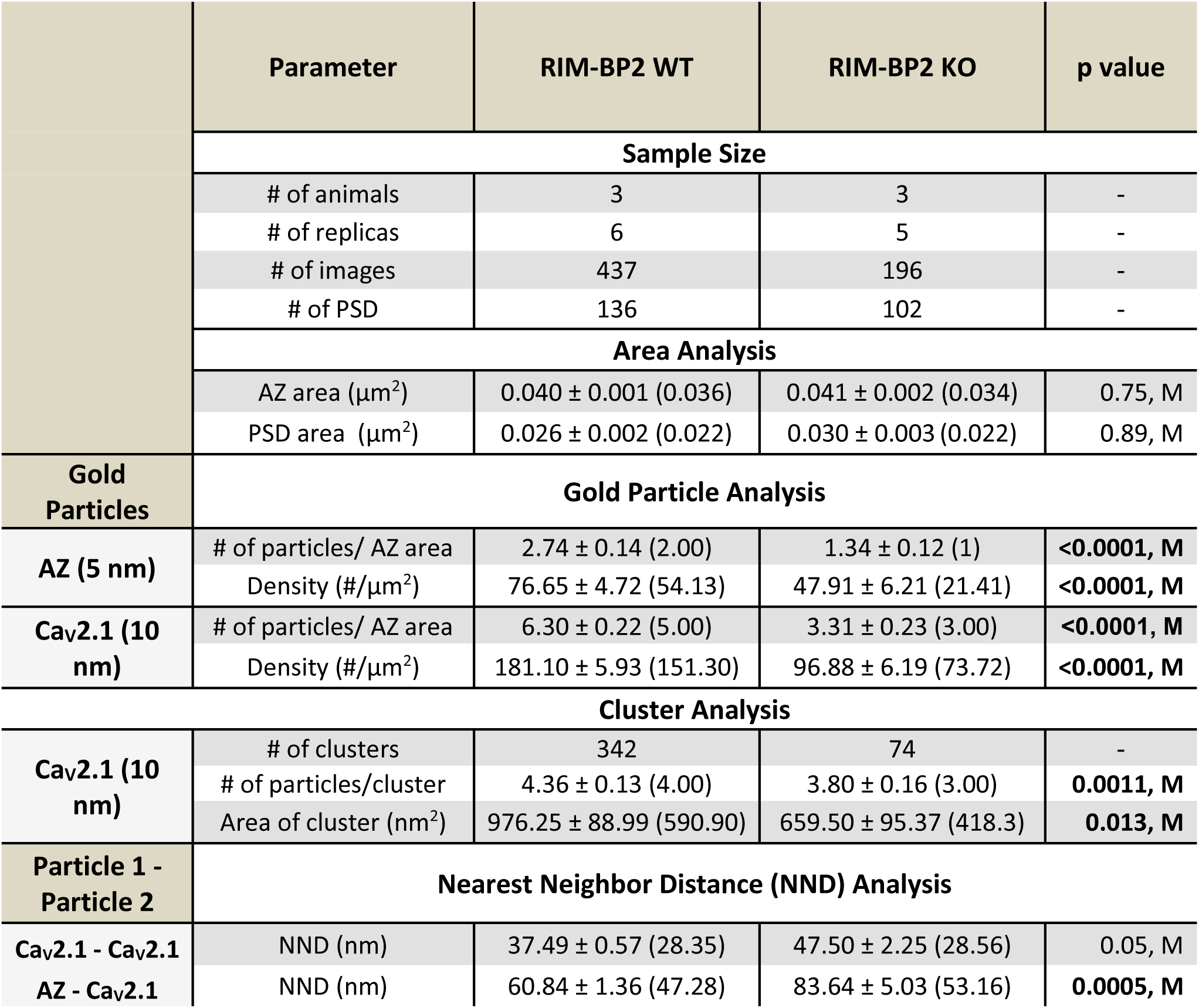
Quantitative analysis SDS-FRIL electron micrographs. Data were distributed non-normally as determined by Kolmogorov-Smirnov test. Statistical significance of the comparison between RIM-BP2 WT and KO was determined by Mann-Whitney U-Test (denoted in the table as M).

A low labelling efficiency for AZ proteins would overestimate the AZ to AZ and Ca_V_2.1 to AZ particles nearest neighbour distances (NNDs). Hence, we only compared the NND for Ca_V_2.1 to Ca_V_2.1 and AZ to Ca_V_2.1 particles. The value of mean + 2 standard deviations of the Gaussian fit to the distribution of Ca_V_2.1 to Ca_V_2.1 NNDs was considered the threshold for the maximum distance by which two particles can be separated and still belong to the same cluster (Fig 9A-B, see materials and methods): mean + 2 s.d. were 43.09 nm and 43.92 nm for WT and RIM-BP2 KO respectively. The threshold used for defining a cluster in our analysis was 40 nm as used in previous analysis (Miki et al., 2017). In many KO AZ images, there were not even three Ca_V_2.1 gold particles to qualify as a cluster (Fig. 9-2C). Within the qualified clusters, we still found a significantly reduced number of Ca_V_2.1 particles per cluster in the RIM-BP2 KO (Table 5) which is consistent with a more dispersed Ca_V_2.1 topography at the AZ as indicated in the STED analysis. To confirm that the clusters obtained in our analysis were valid and not reflecting chance occurrence, we compared our ‘real’ data to 500 random simulations of cluster arrangements (see materials and methods, and (Luján et al., 2018; Kleindienst et al., 2020)). Table 6 confirms the validity of Ca_V_2.1 clusters and NNDs between Ca_V_2.1 to Ca_V_2.1 and AZ to Ca_V_2.1 particles. The major findings of our FRIL analysis are the significantly lower number and density of Ca_V_2.1 channels as well as the increased AZ to Ca_V_2.1 NND in RIM-BP2 KO AZs (Fig. 9C, Table 5). There was a trend toward larger Ca_V_2.1 to Ca_V_2.1 NND at RIM-BP2 KO AZs which did not reach significance (Fig. 9D). The increased AZ to Ca_V_2.1 NND implies a looser coupling between the Ca^2+^ channels and the AZ proteins likely marking SV release sites.

**Table 6.** Comparison of Real and simulated distribution of gold particles. Real data were compared with 500 random and fitted (only NND analysis) simulations to confirm that the observed clusters and NND distribution are significantly different from chance (random) occurrence.

| Genotype | Au Particles | Parameter | Real Data | Simulated Data | p value |
| --- | --- | --- | --- | --- | --- |
|  |  | Sample Size |  |  |  |
|  |  | # of animals | 3 | 3 | - |
|  |  | # of replicas | 6 | 5 | - |
|  |  | # of images | 437 | 196 | - |
| Cluster Analysis |  |  |  |  |  |
| RIM-BP2<br>KO | AZ<br>(5 nm) | # of clusters/AZ | 0.08 ± 0.02 | 0.05 ± 0.01 | 0.015 |
|  |  | # of particles/cluster | 4.33 ± 0.43 | 3.71 ± 0.33 | 0.006 |
|  |  | Area of cluster (nm²) | 759.82 ± 186.11 | 542.89 ± 125.05 | 0.09 |
|  | Ca<sub>v</sub>2.1<br>(10 nm) | # of clusters/AZ | 0.38 ± 0.04 | 0.15 ± 0.02 | 6.13 e-13 |
|  |  | # of particles/cluster | 3.80 ± 0.16 | 3.35 ± 0.06 | 0.0003 |
|  |  | Area of cluster (nm²) | 659.50 ± 95.37 | 438.90 ± 29.90 | 0.005 |
| Particle 1<br>- Particle<br>2 | Particle 1 -<br>Particle 2 | Nearest Neighbor Distance (NND) Analysis |  |  |  |
| WT | Ca<sub>v</sub>2.1 - Ca<sub>v</sub>2.1 | NND (nm) | 42.13 ± 1.29 | 55.20 ± 1.05 | 5.87 e-28 |
| RIM-BP2<br>KO | Ca<sub>v</sub>2.1- Ca<sub>v</sub>2.1 | NND (nm) | 58.14 ± 5.25 | 85.91 ± 4.80 | 1.84 e-06 |

### RIM-BP2 disruption alters the SV organization at the endbulb active zones

Next, we high pressure froze aVCNs slices, acutely prepared as for physiology in order to best relate structure and function, for electron tomography analysis of SV organization at the endbulb AZ. Following freeze-substitution, embedding, sectioning and tomography, we rigorously analysed and reconstructed AZ of RIM-BP2 deficient and WT endbulb synapses (Fig. 10). The AZ area, approximated from the extent of the postsynaptic density, was significantly larger in RIM-BP2-deficient endbulbs (Fig. 10C & Table 7, *p < 0.01*, Student’s t-test). The number (Fig. 10D) and density (Fig. 10E) of SVs per AZ were not altered. For a more comprehensive analysis we compared the SV distribution within 200 nm of the presynaptic AZ membrane (perpendicular to the presynaptic membrane into the cytosol of the presynaptic terminal) in 20 nm bins. Morphologically docked SVs (0-2 nm distance), analyzed separately, were significantly fewer in number at RIM-BP2 KO AZs. Significantly fewer SVs were also found for non-docked membrane proximal SVs (2 to 20 nm) and SVs within 40 to 60 nm from the AZ membrane (Fig 10F, *p < 0.01*, Wilcoxon rank test, Table 7). We also analysed top-views of the AZs showing only the docked SVs in the models generated from the tomograms and quantified the percentage of AZs with zero to eight docked SVs. We found that 50% of the analysed RIM-BP2 KO AZs showed zero docked SVs, while two docked SVs per AZ were most frequently encountered in WT (27% of the AZs, Fig. 10G). We further tested for effects of RIM-BP2 disruption on the lateral distribution of the docked SVs by setting a central point within the generated top views and quantifying the distances of all docked SVs from the center (we note that the AZ area captured in the tomograms might not necessarily allow proper definition of the AZ center). The docked SVs appeared to be further away from the center in RIM-BP2 KO synapses, possibly representing a broader distribution over the whole AZs area (Fig. 10J). To analyse the membrane proximal SVs in more detail, we quantified the proportion of SVs in the 2-20 nm bin and observed a shift towards fewer SVs at mutant AZs. Whereas most of the WT (15%) AZs contained 6 SVs within 2-20 nm from the AZ membrane, most mutant AZs showed only 2 or 5 SVs within this distance (Fig. 10K). Lastly, by measuring the diameter of SVs, we found that they exhibited comparable sizes at RIM-BP2 KO and RIM-BP2 WT endbulb AZs (WT: 51.94 ± 0.55 nm; KO: 52.67 ± 0.54 nm). We conclude that RIM-BP2 contributes to normal SV docking and SV organization in close proximity to the membrane of the presynaptic AZ. These changes seem compatible with the lower number of super-primed SVs, impaired SV replenishment (Figs. 3 and 7) and the reduced release probability (Fig. 3).

**Figure 10.**
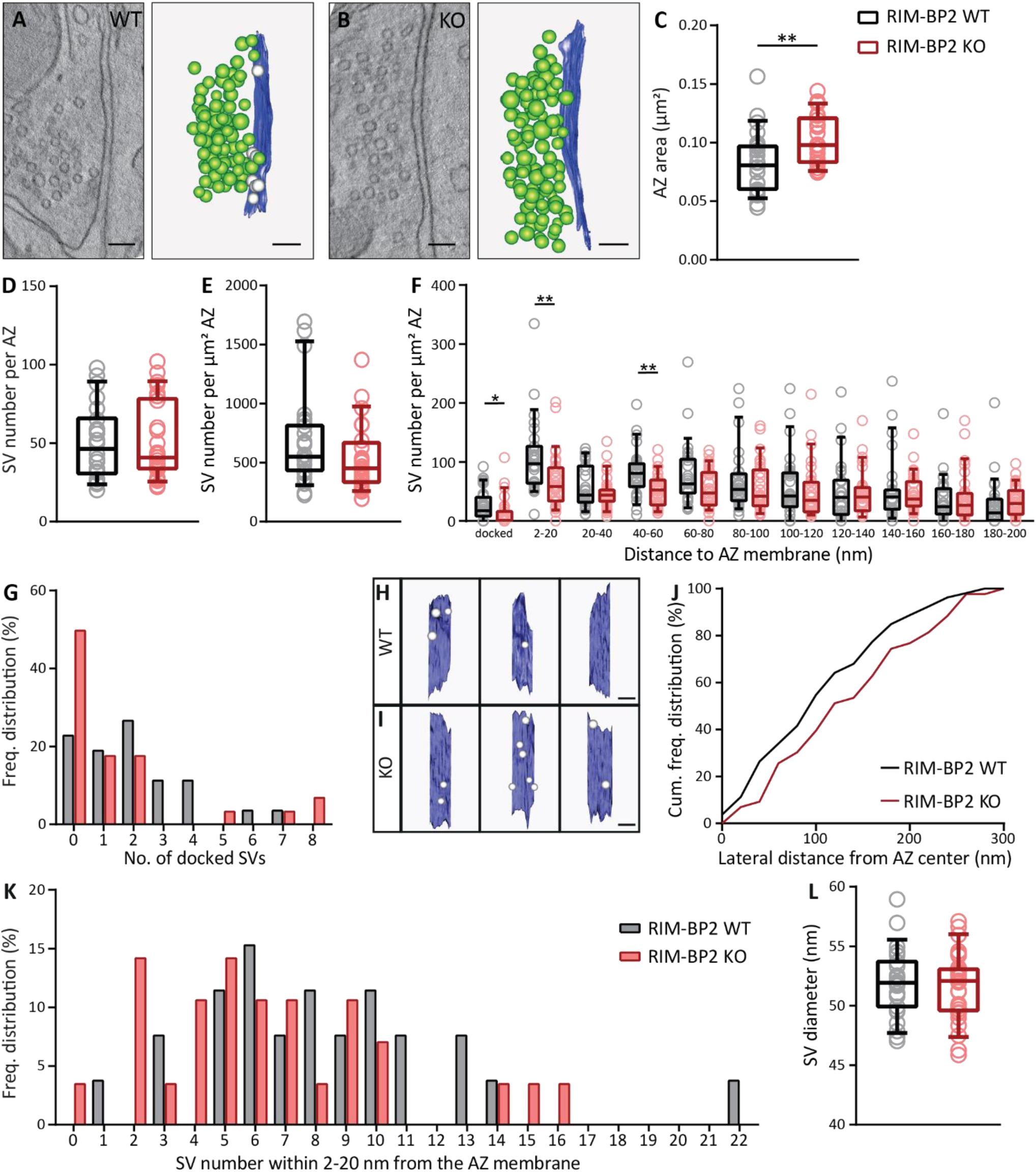
RIM-BP2 disruption alter the axial SV distribution at the endbulb active zones. (**A, B**) Single virtual sections and corresponding models of representative tomograms of RIM-BP2 WT (**A**) and KO (**B**) active zones (AZs) showing the AZ membrane (blue), synaptic vesicles (SVs) (green), and morphologically docked SVs (gray). Scale bars: 100 nm. (**C**) The AZ area estimated by the PSD, was significantly larger in the KO endbulb synapses. \*\**p-value* < 0.01, Student‘s t test. Each data point represents the AZ area of individual synapses. (**D**) Unaltered total number of SVs in mutant AZs. *p-value* > 0.05, Student‘s t test. Each data point represents the number of SVs per AZ of individual synapses. (**E**) SV number normalized to the AZ area is unaltered in KO endbulb AZs. *p-value* > 0.05, Wilcoxon rank test. Each data point represents the number of SVs normalized to the AZ area of individual synapses. (**F**) The number of morphologically docked SVs (0-2 nm) and SVs within 200 nm from the AZ membrane normalized to the AZ area divided into 20 nm bins. \**p-value* < 0.05, \*\**p-value* < 0.01, Wilcoxon rank test. Each data point represents the number of SVs in each bin normalized to the AZ area of individual AZs. (**G**) Frequency distribution of the number of morphologically docked SVs. (**H,I**) Top views of representative tomogram models of RIM-BP2 WT (**H**) and KO (**I**) AZs with docked SVs. Scale bars: 100 nm. (**J**) Cumulative distribution of the lateral distances of morphologically docked SVs to the assumed center of the reconstructed AZ. (**K**) Frequency distribution of the SV number within 2-20 nm from the AZ membrane. (**L**) Mean SV diameter is unaltered in mutant synapses. *p-value* > 0.05, Student‘s t test. Each data point represents the mean diameter of SVs of individual synapses. Box and whisker plots present median, lower/upper quartiles, 10–90^th^ percentiles. RIM-BP2-WT (N = 4; n = 26) in black and RIM-BP2-KO (N = 3; n = 28) in red (N, number of animals; n, number of AZs). For details about mean ± SEM, median, and statistics see Table 7.

**Table 7.**
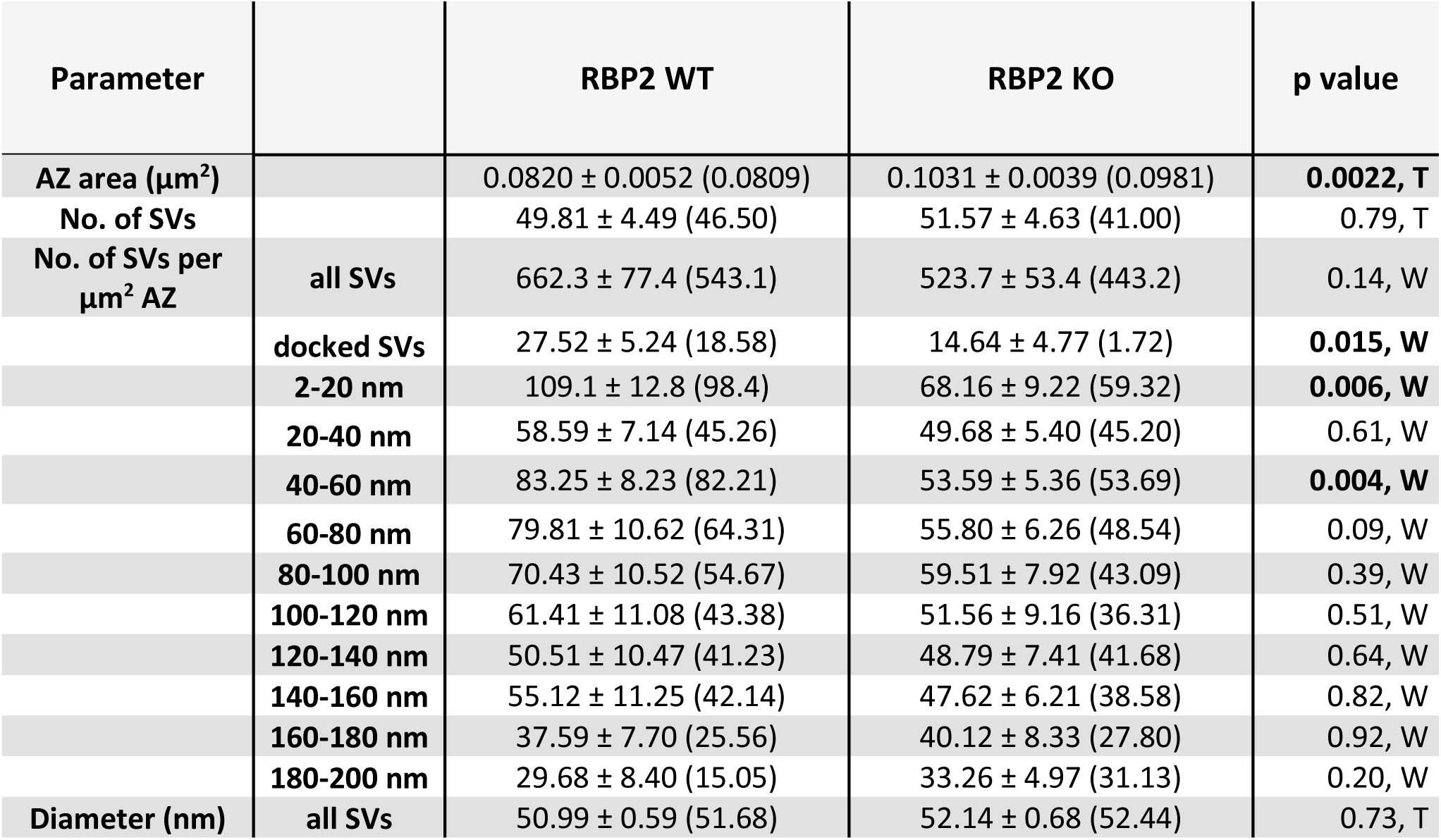
Quantitative analysis of electron tomograms. Data are presented as mean ± SEM and medians are shown in parentheses. Normal distribution was assessed with the Jarque-Bera test and equality of variances was assessed with the F-test in normally distributed data. Statistical significance of normally distributed data was determined by unpaired Student’s t-test (denoted as T), while that of the non-normally distributed data was determined by the Wilcoxon Rank test (denoted as W). n.s.: not significant, * *p-value* < 0.05, ** *p-value* < 0.01.

## Discussion

Priming of SVs, Ca_V_ function as well as the topography of Ca_V_ and SV release sites at the AZ, co-determine the probability of SV release in response to the action potential invading the presynaptic terminal. Here, we probed the role of RIM-BP2, thought to serve as molecular linker between Ca_V_ and release sites, and alternatively in SV priming via Munc13, in synaptic transmission at the endbulb of Held synapse. Using super-resolution immunofluorescence and immuno-electron microscopy we demonstrate that RIM-BP2 disruption alters the topography of Ca_V_2.1 channels at the AZs. Electron tomography revealed fewer docked and membrane-proximal SVs at the AZ. As a physiological corollary of these structural changes, we found a reduction in the amplitude of evoked EPSCs, reduced release probability, and slowed SV replenishment to the RRP. We postulate that RIM-BP2, likely via interaction with Munc13-1, promotes a ‘superprimed’ (Taschenberger et al., 2016) or ‘tightly docked’ (Neher and Brose, 2018) SV state. Moreover, RIM-BP2 organizes the topography of Ca_V_2.1 channels, tightly coupling them to SV release sites.

Synaptic transmission at calyceal synapses of the lower auditory pathway shows impressive temporal fidelity. At the first central relay of the auditory pathway, co-incident transmission from endbulbs formed by ANFs drives the postsynaptic BCs at hundreds of Hz and with microsecond precision as required for time-critical neural computations such as in sound localization (Trussell, 1999; von Gersdorff and Borst, 2002). Such fidelity is enabled by synergistic adaptations on molecular, synaptic and network levels. In the mouse AVCN, BCs receive input from on average 3-4 endbulbs (Cao and Oertel, 2010; Butola et al., 2017) with approximately 400 excitatory AZs (Nicol and Walmsley, 2002; Mendoza Schulz et al., 2014; Butola et al., 2017). Endbulbs feature a large RRP (Lin et al., 2011) and high release probability (estimates range from 0.2 to 0.7: Oleskevich and Walmsley, 2002; Wang and Manis, 2005; Chanda and Xu-Friedman, 2010; Mendoza Schulz et al., 2014; Butola et al., 2017) which enable massive synchronous parallel release for powerful excitation of BCs. RIM-BP2 deletion hampers this reliable and temporally precise transmission of auditory information: firing at sound onset was impaired both in rate and temporal precision (Fig. 5). We mainly attribute the reduced firing at sound onset (40% reduction) to the observed impairment of endbulb transmission (EPSC_1_ amplitude reduction: 36%, Fig. 1), as the deficit in sound onset coding in RIM-BP2 KO ANF was mild (approximately 10% reduction in peak onset firing rate, (Krinner et al., 2017)) and convergence of ANF inputs to BCs is expected to alleviate consequences of impaired ANF coding for BC firing (Joris et al., 1994; Buran et al., 2010). These results from single-neuron recordings are in good agreement with previously reported auditory brainstem responses (Krinner et al., 2017), in which the amplitude reduction was lower for wave I (approximately 30 %) than for wave III (approximately 70%). Wave I and III are attributed to the synchronized firing of ANFs and cochlear nucleus neurons respectively (Melcher et al., 1996). Future studies, also employing analysis of gap detection or sound localization should evaluate the behavioural consequences of this temporal processing deficit.

Impaired transmission of sound onset information at RIM-BP2-deficient endbulbs is primarily rooted in a nearly halved initial release probability, which could be restored to wildtype levels upon increasing Ca^2+^ influx (Fig. 7). We attribute the decreased release probability to i) the altered topography of Ca_V_2.1 and SVs and ii) to the impaired SV docking at RIM-BP2-deficient endbulb AZs. Whole-cell patch-clamp recordings showed normal Ca^2+^ currents in RIM-BP2-deficient endbulbs (Fig. 2) and confocal imaging of Ca_V_2.1 immunofluorescence semi-quantitatively reported a normal Ca^2+^ channel complement of the AZs (Fig. 8-1). This suggests that unlike ribbon synapses (Krinner et al., 2017; Luo et al., 2017), RIM-BP2 is not strictly required for establishing a normal presynaptic Ca_V_ complement. However, both FRIL and STED imaging of immunolabeled Ca_V_2.1 channels revealed an impaired clustering of Ca_V_2.1 at RIM-BP2-deficient endbulb AZs. The number and density of Ca_V_2.1 immunogold particles was reduced at the AZs and the nearest neighboring Ca_V_2.1 was further away from an immunogold particle marking an AZ protein (Table 5). Nonetheless, distribution of the Ca_V_2.1 immunogold particles was significantly different from random, likely reflecting the presynaptic Ca_V_ organization by other multi-domain proteins of the AZ such as RIMs (Han et al., 2011; Kaeser et al., 2011; Jung et al., 2015) and CAST (Dong et al., 2018; Hagiwara et al., 2018). Immunofluorescence of Ca_V_2.1 as well as of bassoon (marking the presynaptic density) was less confined in space and immunofluorescent spots were more oval in shape in the absence of RIM-BP2 as compared to compact, round spots in the WT (Figs. 8 and 8-2). Moreover, the nearest neighboring immunofluorescence spot was further away both for Ca_V_2.1 and bassoon. In summary, our data indicate that RIM-BP2 contributes to orchestrating Ca_V_2.1 channels at the AZ. Based on our data and in line with previous studies on other synapses (Liu et al., 2011; Acuna et al., 2015; Brockmann et al., 2020; Petzoldt et al., 2020), we hypothesize that RIM-BP2 via its interaction with Ca_V_2.1 and AZ proteins, contributes to tight coupling of Ca^2+^ channels and vesicular release sites also at the endbulb of Held synapse. Future experimental and theoretical studies will be needed to further test this hypothesis.

Recently, an alternative interaction of RIM-BP2 with the C2B domain of Munc13-1 has been reported by which RIM-BP promotes release probability via SV docking/priming (Brockmann et al., 2020). The Munc13-1 C2B domain carrying the KW mutation (Shin et al., 2010), showed the highest affinity for RIM-BP2 binding. Of note, we did not observe a reduction in Munc13-1 or in any other major AZ protein in our semi-quantitative analysis of AZs in RIM-BP2 deficient endbulbs of Held (Fig. 6). This provides confidence in attributing functional and morphological alterations to the lack of RIM-BP2 function rather than to quantitative changes in other AZ proteins. Our functional and electron tomographic analysis of SV pool organization at RIM-BP2-deficient endbulbs now provides evidence for a role of RIM-BP in SV priming, likely via its interaction with Munc13-1. Rigorous electron tomography analysis of docking showed nearly halved numbers of morphologically docked SVs (defined in a distance from 0-2 nm to the AZ membrane) at RIM-BP2-deficient AZs, which might represent a ‘tight docking’ (Neher and Brose, 2018). We speculate this to indicate that Munc13-1-mediated docking that generates primed SVs (Siksou et al., 2009; Imig et al., 2014) partially depends on the interaction with RIM-BP2. The reduction of both docked SVs and those at a distance of 2-20 nm (membrane proximal SVs) from the plasma membrane might then suggest that the upstream loose-docking is rate limiting or less stable in the absence of RIM-BP2. An alternative explanation for retarded recovery from depression might be that RIM-BP2 facilitates release site clearance (Neher and Sakaba, 2008).

Yet, our analysis of pool dynamics in regular [Ca^2+^]_e_ indicated an unaltered RRP size despite the fact that the numbers of docked and membrane-proximal SVs were nearly halved. Interestingly, we found a trend toward a smaller RRP in RIM-BP2-deficient endbulbs when restoring release probability by enhanced Ca^2+^ influx. Despite this partial restoration, the first evoked EPSC amplitude in the absence of RIM-BP2 was still only 50% of that in the WT synapses (Fig. 7, table 4). This persisting difference alludes to a deficit in ‘super-primed’ (Fig. 3, table 2) or ‘tightly docked’ SVs (Fig. 10, table 7). It is tempting to speculate that the impaired SV docking is uncovered under conditions that occlude the effect of altered spatial Ca^2+^ channel-release site coupling on release probability. Our morphological analysis of Ca^2+^ channel-release site coupling is hampered by i) the low number of docked SVs in electron tomography and ii) the lack of information on the Ca_V_ position in electron tomography and of SV docking in FRIL electron micrographs. Nonetheless, there was a trend for docked SVs to be further away from the estimated AZ center, possibly reflecting a more random SV topography due to lack of RIM-BP2-mediated interaction with Ca_V_s. In addition, or alternatively, the RRP estimated by SMN/EQ analyses of responses to train stimulation might contain SVs that undergo tethering, docking and fusion during the train. Recent work by (Pofantis et al., 2021), showed that deleting the presynaptic protein Mover specifically affects the high P_r_ component of the RRP at the Calyx of Held. Pofantis et al. analysed eEPSC trains with non-negative matrix factorization to reveal components representing sub-pools of SVs with different contributions to transmitter release during the train. Future studies that tackle how such sub-pools deplete and how they recover from depletion in synapses lacking both RIM-BP2 as well as RIM-BP2 and Mover will help to further dissect the contributions of RIM-BP2 to SV super-priming and Ca_V_ clustering as determinants of P_r_.

Exciting topics for future studies include i) the relative contributions of altered Ca^2+^ channel-release site coupling and impaired SV docking to the reduced release probability, ii) a potential contribution of release site clearance to the observed deficit in SV docking, and iii) the molecular mechanisms and structure-function relationship of SV replenishment. Our morphological and functional experiments indicate that RIM-BP2 takes a role in tethering SVs near the plasma membrane *en route* to docking. Functionally, the fast, Ca^2+^ dependent component of recovery from depression due to trains stimulation was hampered. This could reflect a lower local [Ca^2+^] due to mislocalization of Ca_V_2.1 channels. Indeed, a key role of Ca^2+^ in regulating SV replenishment at calyceal synapses has been reported in multiple studies (Wang and Kaczmarek, 1998; Hosoi et al., 2007). Interestingly, enhancing Ca^2+^ influx by elevated [Ca^2+^]_e_ uncovered a reduced SV replenishment during train stimulation, which would seem reflect a Ca^2+^ independent limitation e.g. of SV priming. Future studies on calyceal synapses of mice carrying mutations that target Ca^2+^ dependent effects on Munc13-1 function and/or the RIM-BP2-Munc13-1 interactions by combined functional and ultrastructural analyses will be required to elucidate this intricate process. To best relate AZ structure and function, future studies will ideally employ optogenetic or electric field stimulation followed by high pressure freezing and EM tomography (Watanabe et al., 2013; Imig et al., 2020 p.202).

## Author Contribution

This study was conceived by T.M. and T.B. The experimental work was performed by T.B. (slice electrophysiology, *in vivo* electrophysiology, electron microscopy in the lab of R.S. with help of P.K., D.K. and R.S.), T.A. (slice electrophysiology, immunohistochemistry) and A.H. (electron microscopy). T.M., T. A., and T.B. prepared the manuscript with contributions of A.H. and C.W.

## Acknowledgements

We thank S. Gerke, A.J. Goldak and C. Senger-Freitag for expert technical assistance, G. Hoch for developing image analysis routines, and S. Chepurwar and N. Strenzke for technical support and discussion regarding *in vivo* experiments. We thank Drs. Christian Rosenmund, Katharina Grauel and Stephan Sigrist for providing RIM-BP2 KO mice. We thank J. Neef for help with the STED imaging and image analysis. We thank E. Neher and S. Rizzoli for discussion and comments on the manuscript. Our gratitude also goes to C. H. Huang and J. Neef for constant support and scientific discussion. This work was funded by the Deutsche Forschungsgemeinschaft (DFG, German Research Foundation) through the Collaborative Sensory Research Center 1286 (to C.W. (A4) and T.M. (B5)) and under Germany’s Excellence Strategy - EXC 2067/1-390729940.

